# Working memory signals in early visual cortex do not depend on visual imagery

**DOI:** 10.1101/2023.02.13.528298

**Authors:** Simon Weber, Thomas Christophel, Kai Görgen, Joram Soch, John-Dylan Haynes

## Abstract

It has been suggested that visual images are memorized across brief periods of time by vividly imagining them as if they still were there. In line with this, the contents of both working memory and visual imagery are known to be encoded already in early visual cortex. If these signals in early visual areas were indeed to reflect a combined imagery and memory code, one would predict them to be weaker for individuals with reduced visual imagery vividness. Here, we systematically investigated this question in two groups of participants. Strong and weak imagers were asked to remember images across brief delay periods. We were able to reliably reconstruct the memorized stimuli from early visual cortex during the delay. Importantly, in contrast to the prediction, the quality of reconstruction was equally accurate for both strong and weak imagers. The decodable information also closely reflected behavioral precision in both groups, suggesting it could contribute to behavioral performance, even in the extreme case of completely aphantasic individuals. Our data thus suggest that working memory signals in early visual cortex can be present even in the (near) absence of phenomenal imagery.

## Introduction

In recent years, visual imagery, the ability to generate pictorial mental representations in the absence of external visual stimulation (Kosslyn & Thompson, 2003; Pearson & Kosslyn, 2015), has received increasing attention as a potential mechanism for supporting visual working memory (Albers et al., 2013; Tong, 2013).

Both visual imagery and visual working memory have been linked to the encoding of information in early visual cortex (Dijkstra et al., 2019; Klein et al., 2004; Kosslyn & Thompson, 2003; Lee & Baker, 2016; Serences, 2016). The sensory recruitment hypothesis of visual working memory (D’Esposito & Postle, 2015; Sreenivasan et al., 2014) posits that visual information is maintained using selective activation patterns in early visual cortex. This matches with a common view of visual imagery, where early visual areas encode detailed, perception-like mental images via top-down connections from high-level regions (Dijkstra et al., 2017; Mechelli, 2004). Encoding of contents has been reported to be similar between perception and visual working memory (Ester et al., 2009; Harrison & Tong, 2009; Lee et al., 2013; Serences et al., 2009). This similarity has also been shown to hold between perception and imagery across multiple features, including orientations (Albers et al., 2013), objects (Cichy et al., 2012; Lee et al., 2012; Ragni et al., 2020; Reddy et al., 2010), letters (Senden et al., 2019), and natural scenes (Naselaris et al., 2015). Furthermore, both visual working memory (Teng & Kravitz, 2019) and visual imagery (Pearson et al., 2008) can interfere with and bias perception of subsequent stimuli.

The similarities in cortical organization of imagery and visual working memory raise the question whether these two processes might be related or even share the same neural substrate. Indeed, it was directly shown for normal-viewing participants that visual working memory and imagery representations of orientations exhibit very similar neuronal activity patterns in early visual cortex (Albers et al., 2013) suggesting that visual working memory and visual imagery share a similar neural substrate (Tong, 2013). In this view, participants might briefly memorize visual stimuli in working memory tasks by vividly imagining them across the delay period.

However, the ability to generate imagery as well as its vividness differ substantially across individuals (Kosslyn et al., 2001). Some people even report the complete absence of phenomenal imagery (“aphantasia”; Zeman et al., 2015; Zeman et al., 2010). Nonetheless, these differences do not appear to manifest themselves systematically in behavioral measures of memory. Rather, most studies indicate that behavioral performance in visual working memory tasks is comparable across imagery vividness levels, including the extreme case of aphantasic individuals (Jacobs et al., 2018; Zeman et al., 2015). However, differences have been reported. For example, working memory performance for strong imagers is disrupted by irrelevant visual input, while weak imagers show no such distraction effect (Keogh & Pearson, 2014), indicating the use of distinct memorization strategies. This is supported by comparing reports of strong and weak imagers. Strong imagers report to rely mostly on visual strategies when solving visual working memory tasks. In contrast, weak imagers tend to report using different cognitive strategies such as verbal or categorical associations (Bainbridge et al., 2021; Keogh et al., 2021; Logie et al., 2011). Thus, visual imagery might to be only one of several cognitive tools that can be used to solve visual working memory tasks. If this is true, then weak imagers could use different representational systems for maintaining stimulus features other than sensory recruitment in early visual cortex.

In line with this, the cognitive-strategies framework of working memory (Pearson & Keogh, 2019) postulates that the cognitive strategy used to solve a working memory task determines the format in which a stimulus is represented in the brain, and consequently influences how much information about the stimulus is present within a given cortical region. In the case of visual imagery, this could mean that individuals with high imagery vividness spontaneously recruit their early visual cortex to maintain detailed stimulus representations, while individuals with low imagery vividness employ alternative, non-visual strategies to solve the same cognitive task. Together, this predicts that strong imagers should retain more information about a stimulus feature in their visual cortex activity than weak imagers.

Here, we directly test this hypothesis by assessing the influence of imagery vividness on the strength of visual working memory representations in visual cortex, using functional magnetic resonance imaging (fMRI). We recruited two groups of study participants, one with very high and one with very low imagery vividness scores as assessed by an established questionnaire (VVIQ, Fig 1B, see Methods; Marks, 1973). In the main experiment, participants performed a working memory task that involved memorizing a bright orientation stimulus across a brief delay (Figure 1A). We used a brain-based decoder (periodic support vector regression; see Methods) to reconstruct these orientations from brain activity patterns in early visual cortex obtained during the memory delay period. If strong imagers indeed rely more on imagery signals in early visual cortex to maintain the stimulus across the delay, this could lead to two predictions: First, that sensory information should be represented more accurately in the early visual brain signals of strong as opposed to weak imagers; second, sensory information in early visual areas should also be more predictive of an individual’s behavioral performance, especially in strong imagers.

**Figure 1.**
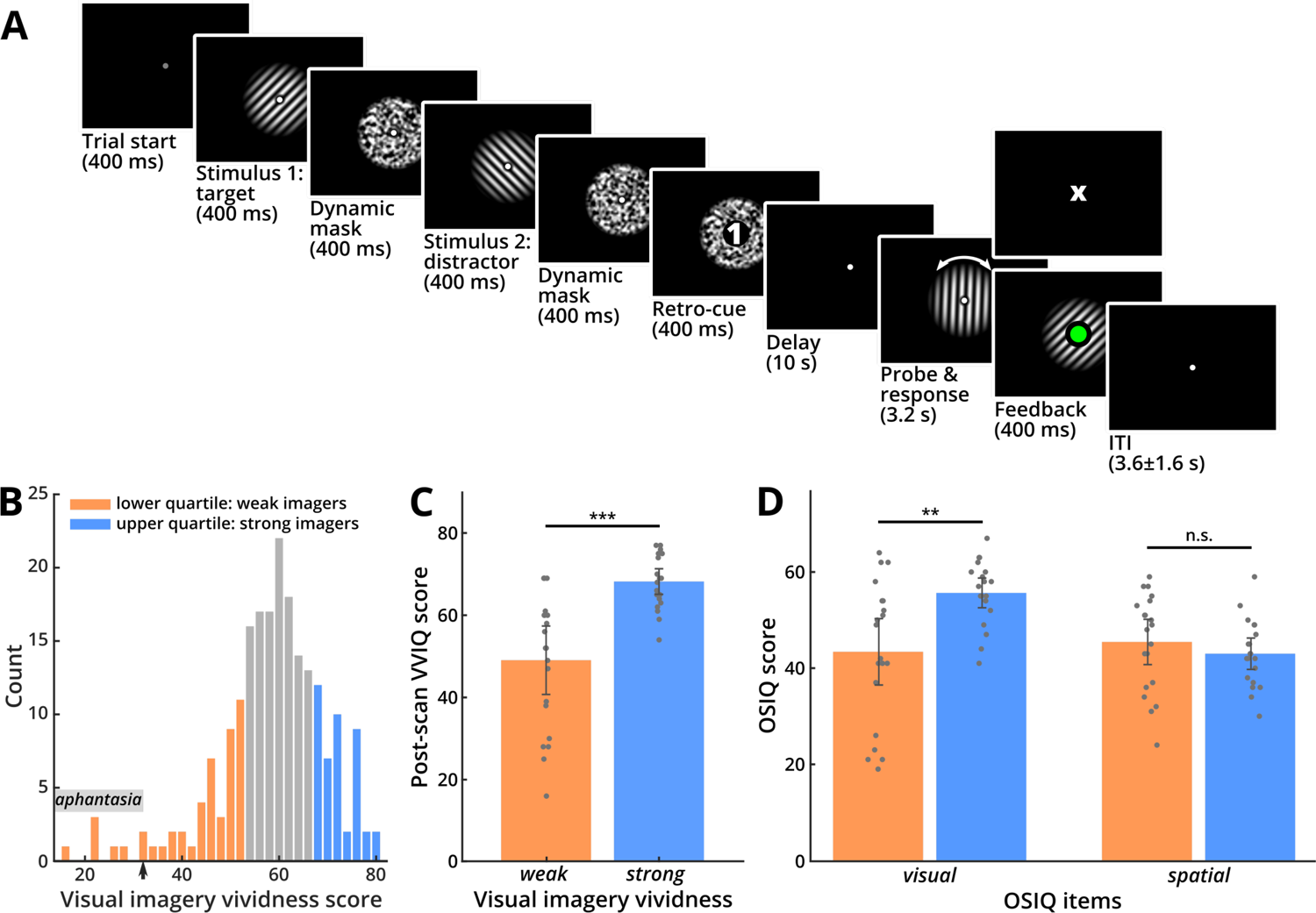
Experimental task and questionnaire data. **(A)** Sequence of events in one trial of the experiment. In each trial, participants were successively presented with two orientation stimuli, each followed by a dynamic noise mask. Orientations were drawn from a set of 40 discrete, equally spaced orientations between 0° and 180°. The stimuli were followed by a numeric retro-cue (“1” or “2”), indicating which one of them was to be used for the subsequent delayed-estimation task (“target”), and which could be dropped from memory (“distractor”). The orientation of the cued target grating had to be maintained for a 10-second delay. After the delay a probe grating appeared, which had to be adjusted using two buttons and then confirmed via an additional button press. Subsequently, visual feedback was given indicating whether a response was given in time (by turning the fixation point green, lower panel) or missed (by displaying a small “X” at the end of the response period if no response was given in time, upper panel). Cue and feedback are enlarged in this illustration for better visibility. **(B)** Distribution of the scores in an online visual imagery questionnaire (VVIQ, see Methods) that was used for recruitment. Subjects from the upper (blue) versus lower (orange) quartiles of the distribution were recruited for the strong and weak imagery vividness groups, respectively. The small arrow on the x-axis points to the aphantasia cutoff. **(C)** Questionnaire scores of the post-scan (repeated) VVIQ for weak and strong imagers, as defined by the recruitment scores. The post-scan scores of the weak imagery group were significantly lower than those for the strong imagery group, indicating that the groups were consistent across the study and repeated testing (*t*(38) = −5.086, *p* < 0.001, two-tailed; error bars: 95 % confidence intervals). **(D)** Results for the visual and spatial items from the OSIQ. Scores for the visual items were significantly lower for weak imagers (*t*(38) = −3.338, *p* = 0.002, two-tailed). Scores for the spatial items did not differ between groups (*t*(38) = 0.895, *p* = 0.377, two-tailed; error bars: 95 % confidence intervals), as expected from previous work (Bainbridge et al., 2021; Keogh & Pearson, 2018).

## Results

### Questionnaire data

Study participants were selected via an online version of the established Vividness of Visual Imagery Questionnaire (VVIQ, 210 respondents, Figure 1B; Marks, 1973). We recruited 20 participants each from the lower and upper quartile of the VVIQ score distribution, resulting in two experimental groups (average VVIQ score; weak: 40.75 ± 11.571; strong: 70.7 ± 3.262). After the second fMRI session, each participant repeated the VVIQ and also completed the Object Spatial Imagery Questionnaire (OSIQ; Blajenkova et al., 2006). VVIQ scores had a high test-retest reliability (r = 0.867, p < 0.001), and thus also the difference between weak and strong imagers, as defined by the recruitment scores, was stable across the study period (Figure 1C; *t*_(38)_ = −5.086, *p* < 0.001, two-tailed). In line with previous studies, the OSIQ scores (Figure 1D) had a significant difference between weak and strong imagers for the visual items (*t*_(38)_ = −3.338, *p* = 0.002, two-tailed), but no such difference for the spatial items (*t*_(38)_ = 0.895, *p* = 0.377, two-tailed). Crucially, this pattern of OSIQ results replicates earlier findings obtained with this scale for weak and strong imagers (Bainbridge et al., 2021; Keogh & Pearson, 2018), which serves as a validation of the VVIQ scores as a recruitment measure.

### Behavioral results

Figure 2A shows how accurately participants performed in the task. The figure plots the deviation between participants’ judgements and the true orientations for each trial (grey bars), revealing that the responses were highly accurate. To assess this quantitatively, we fitted a computational model to the response distribution of each participant that yields estimates for behavioral precision and bias (von Mises mixture model; Figure 2A, black line; see Methods for details). Across all participants, responses were precise (precision *K*_1_ = 5.673 ± 2.377), with a small but significant bias to respond anti-clockwise of the target (*μ* = −0.889° ± 1.635°; Figure 2A, inset).

**Figure 2.**
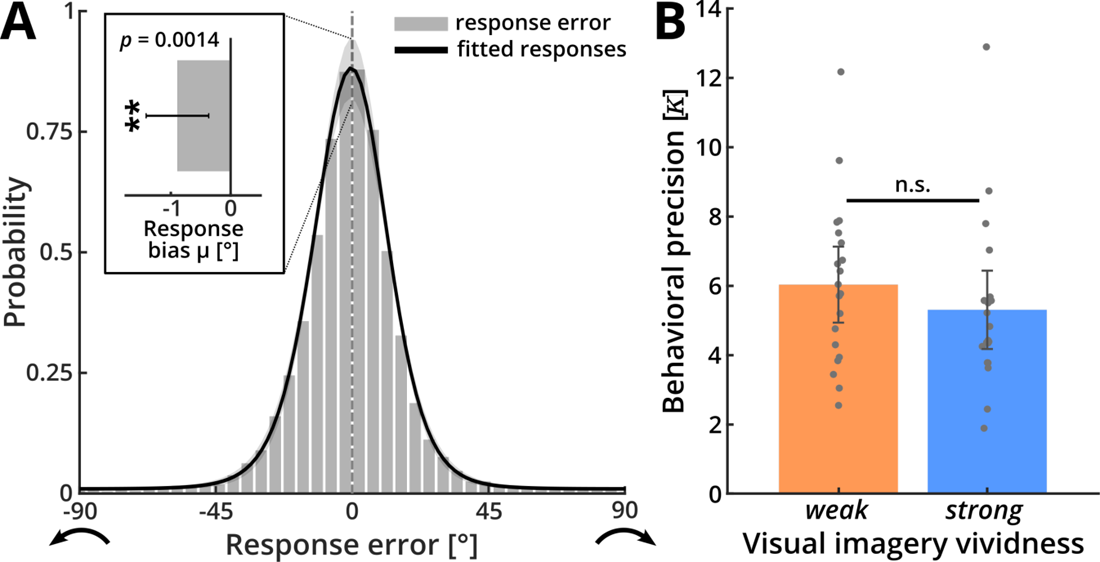
Behavioral results. **(A)** Histogram of deviations between the reported and the true orientation of the target stimuli (grey bars) and a model fit of behavioral responses across all subjects (black line). For this, the responses were modeled using a von Mises mixture model (vMMM) for detections (responses to target orientations, assumed to follow a von Mises distribution with mean 0° plus bias *µ* and behavioral precision *κ*1), swap errors (false responses to distractor orientations, following the same assumptions as detections) and guesses (assumed to follow a continuous uniform distribution between −90° and +90°). The model estimated individual probabilities for each of these three event classes (resulting in mixture coefficients, *r1, r2* and *r3*, respectively). The estimated parameters indicate that participants accurately performed the task: they correctly responded to the target direction in around 95 % of trials (*r*1 = 0.947 ± 0.063). Across participants, responses were precise (*κ*1 = 5.673 ± 2.377), with a small but significant bias to respond anti-clockwise of the target (inset; *μ* = −0.889 ± 1.635°; *t*(39) = −3.437, *p* = 0.0014, two-tailed; error bar: 95 % confidence interval). See Figure S1 for details on the other estimated parameters. **(B)** Behavioral precision (*κ*1) for strong and weak imagers separately. Behavioral precision did not significantly differ between groups (error bars: 95 % confidence intervals).

Importantly, there were no significant differences between strong and weak imagers for behavioral precision (Figure 2B; *t*_(38)_ = −0.965, *p* = 0.341, two-tailed) or any other of the estimated behavioral parameters (Figure S1). This indicates that the high individual differences in visual imagery were not associated with performance differences in the visual working memory task. We used a Bayesian analysis to assess the evidence for absence of a difference in behavioral precision between the weak and strong imagery groups. The Bayes factor indicated that the data were 2.2 times more likely under the null hypothesis (BF_01_ = 2.239) which provides weak evidence for the absence of an effect of imagery vividness on behavioral precision (Jeffreys, 1998).

### Orientation reconstruction from fMRI data

We used a brain-based decoder to reconstruct orientation representations encoded in the patterns of signals in early visual cortex (V1-V3, see Methods). Across all subjects, we were able to reconstruct the true physical target orientation above chance-level for an extended period following delay onset (Figure 3A, green line): At 5 s after delay onset, the accuracy rose to 12 % above chance, where it plateaued until 3 s after probe onset. Following probe onset, the accuracy increased steeply before falling back towards baseline. This later peak in reconstruction performance is likely to reflect the perceptual information of the adjustable probe grating after it had been rotated by the participants to report the target orientation. Reconstruction of the reported orientation yielded a very similar pattern of results (Figure 3A, red line). This close resemblance was expected, given the close match between target and reported orientations (see Figure 2A).

**Figure 3.**
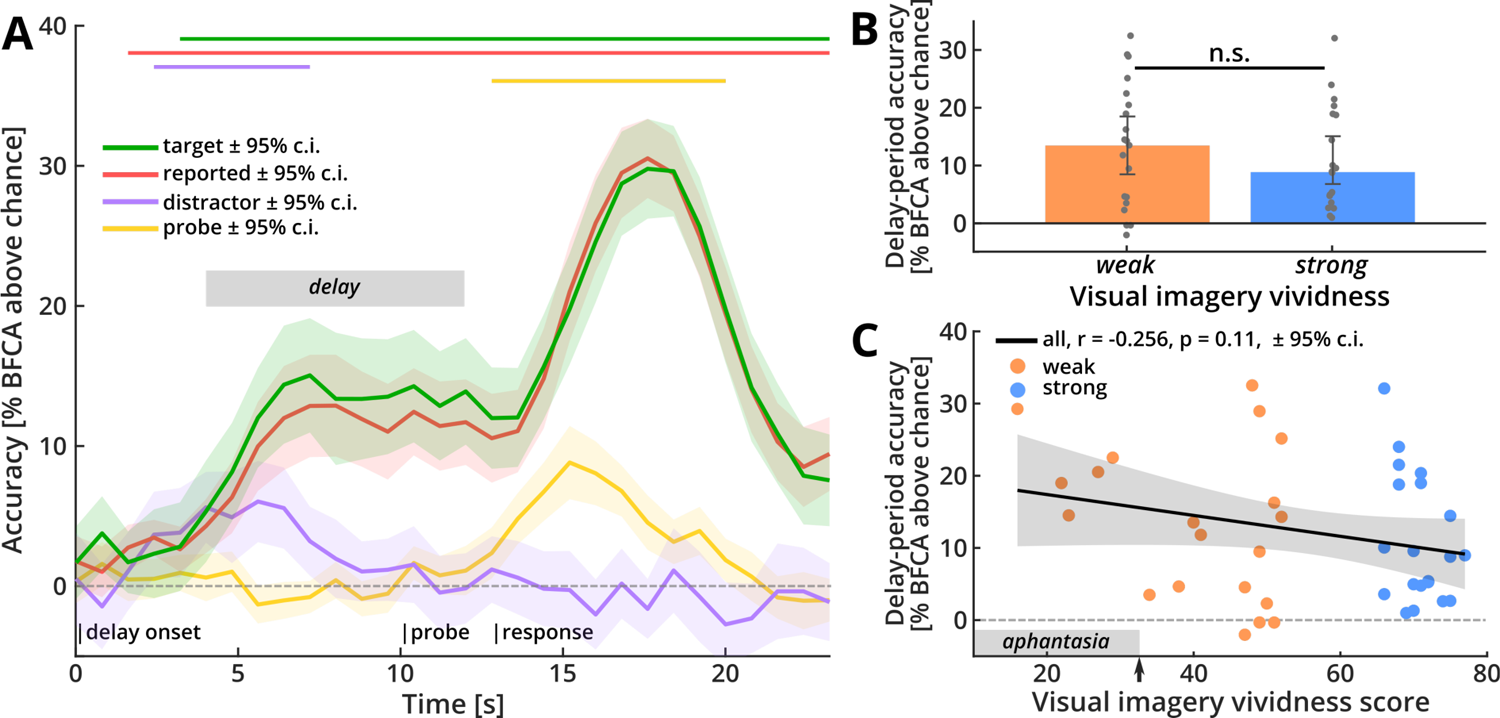
Orientation reconstruction from early visual cortex. **(A)** Reconstruction performance for orientations based on brain signals from early visual areas V1-V3. The y-axis plots the accuracy (BFCA, see Methods), across time for target (green), reported (red), distractor (purple) and probe (yellow) orientations. The horizontal lines above the graph indicate time periods where this reconstruction was significantly above chance (permutation-based cluster-mass statistic, see Methods). The target orientation (green) could be reconstructed above chance-level throughout the delay and report periods (cluster-*p* < 0.001). Reconstruction of the reported orientation (red) followed a highly similar pattern (cluster-*p* < 0.001). The distractor orientation (purple) could only be reconstructed early in the trial (cluster-*p* < 0.001), before falling back to baseline. Reconstruction of the adjustable probe orientation (yellow) was only possible late in the trial (large cluster: cluster-*p* < 0.001; small cluster: cluster-*p* = 0.015), after it had been presented (shaded areas: 95 % confidence intervals). The gray box marks the preregistered delay-period time window used for subsequent analyses. **(B)** Target reconstruction performance for strong and weak imagers separately, pooled across the preregistered delay-period (gray bar in (A)). Delay-period decoding accuracy did not differ between weak and strong imagers (*t*(38) = 0.821, *p* = 0.417, two-tailed; error bars: 95 % confidence intervals). **(C)** Detailed correlation between delay-period accuracy (BFCA) and visual imagery score. There was no significant correlation between the strength of delay-period representations and imagery vividness even when using the full graded imagery scores (shaded area: 95 % confidence interval). Neural information during the delay-period was significantly above chance-level even for aphantasic individuals with a visual imagery score below 32 (grey bar at x-axis; *t*(4) = 8.758, *p* < 0.001, one-tailed, E.A.). The arrow on the x-axis points to the aphantasia cutoff.

We also conducted several checks to test for other predictions of our analysis. First, we reconstructed the orientation of the distractor, i.e., the task-irrelevant orientation stimulus that was not cued and could thus be forgotten after the retro-cue. As expected, information about this distractor orientation (Figure 3A, purple line) was only present briefly at the beginning of the trial after which the accuracy returned to chance-level for the remainder of the trial. In line with previous work on the representation of task-irrelevant stimuli (Albers et al., 2013; Ester et al., 2013; Harrison & Tong, 2009), this transient early information presumably reflects the perceptual signal following the presentation of the distractor early in the trial, delayed by the hemodynamic lag. Second, we reconstructed the initial random starting orientation of the adjustable probe grating (Figure 3A, yellow line). As expected, this resulted in an informative time window late in the trial, after probe onset, likely reflecting the perceptual signal of the adjustable probe before it was rotated for the behavioral response. Together, this pattern of results indicates the presence of sustained, content-selective representations of the memorized stimuli during the delay-period, while task-irrelevant stimulus information was quickly dropped from memory. In an additional analysis we confirmed that the decodable information was not related to systematic eye-movements (Figure S2).

### Group differences in delay-period representations

Next, we proceeded to address the key question whether there was any indication that strong and weak imagers differed in their memory-related information in early visual cortex. Despite robust group-wise reconstruction performance, reconstruction accuracy did not differ between strong and weak imagers (Figure 3B; *t*_(38)_ = 0.821, *p* = 0.417, two-tailed). This was confirmed by a post-hoc Bayesian *t*-test, which provided moderate evidence in favor of the null hypothesis over our original prediction that the early visual cortex signal of strong imagers should contain more information about the stimulus (BF_01_ = 5.275).

To further corroborate the effect, we assessed the possibility that the effect of imagery vividness is more gradual in nature and thus might not be captured by the categorical group difference. To address this, we calculated the correlation between delay-period accuracies and graded imagery vividness scores. Again, the result was not significant (Figure 3C; *r* = −0.256, *p* = 0.11), with strong evidence for the absence of a positive correlation (BF_01_ = 12.442). There was also no relationship between working memory signals and any of the post-scan imagery assessments (see Table S1). Note that delay-period accuracy was significantly greater than chance-level even for the five participants with a visual imagery score of below 32 (marked with a grey bar on the x-axis of Figure 3C; one-sample *t*-test: *t*_(4)_ = 8.758, *p* < 0.001, one-tailed; E.A.), which is generally considered the threshold for aphantasia (Zeman et al., 2015). Taken together, these results suggest that imagery vividness, at least in the form of subjective questionnaire scores, does not affect the strength of delay-period representations of target orientations in early visual cortex.

Finally, we tested a further prediction that would be expected if strong imagers relied more on sensory information encoded in early visual cortex than weak imagers. In that case, there should be a tighter predictive link between behavioral performance and the encoding of information in early visual areas, especially for strong imagers. For this, we assessed whether there was more performance-predictive information in early visual areas of strong imagers. In this additional analysis (E.A.) we observed a strong correlation between delay-period accuracy and behavioral precision (Figure 4A; *r* = 0.728, *p* < 0.001), which was the same across groups (Figure 4B; strong: *r* = 0.81, *p* < 0.001; weak: *r* = 0.657, *p* = 0.002). Interestingly, half of the variance in delay-period accuracy could be explained by behavioral precision (*R^2^*, all: 0.53; strong: 0.656; weak: 0.432). This strong effect suggests that the signals in early visual cortex could potentially play a direct role in maintaining the sensory stimulus across the memory delay (as suggested by the sensory recruitment hypothesis), and that this does not depend on whether a person is a strong or a weak imager.

**Figure 4.**
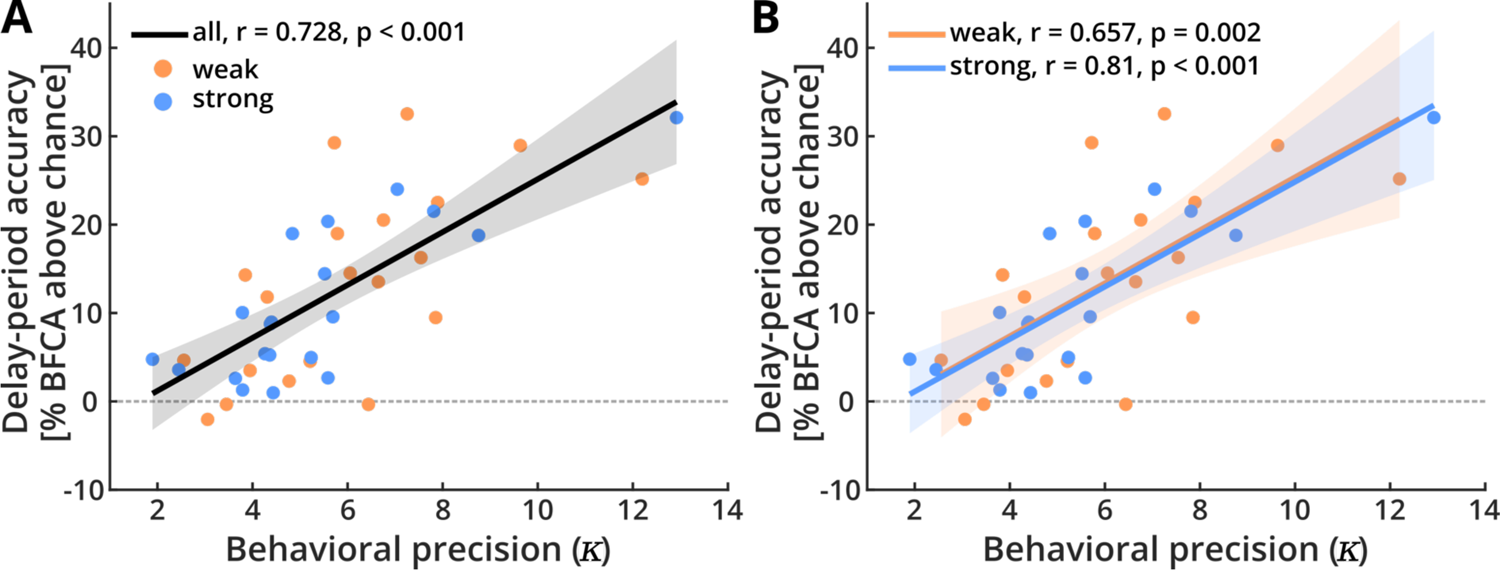
Behavioral precision versus decodable neural information from early visual cortex. Correlation between the behavioral precision (kappa, *K*1) in the task and the accuracy of brain-based reconstruction. The strength of delay-period representations was highly predictable of behavioral precision, both **(A)** across all participants and **(B)** within strong and weak imagery vividness groups. Shaded areas indicate 95 % confidence intervals.

## Discussion

In this study, we investigated to which extent an individual’s visual imagery vividness affects the strength of working memory representations in their visual cortex. Two experimental groups, strong and weak imagers, performed a visual working memory task, which involved memorizing images of oriented lines over a delay. In both groups we found that early visual cortex contained robust information about the remembered orientations across the entire delay period. Importantly, the level of this information did not differ between strong and weak imagery groups. There was also no apparent dependency of visual cortex representations on any other subjective measure of encoding strategy (see Table S1), suggesting that remembered orientations were encoded equally strongly in the visual areas irrespective of an individual’s imagery vividness. Crucially, even the five participants with a VVIQ score of below 32, which is generally considered the threshold for complete absence of phenomenal imagery (“aphantasia”; Zeman et al., 2015) showed comparable visual neural information to the strong imagers (see Figure 2C). Our results therefore show that working memory signals can be present in early visual cortex even in the (near) absence of phenomenal imagery.

While working memory signals in early visual cortex were not modulated by imagery vividness, we did observe a strong correlation between encoded information and individual behavioral precision. Moreover, the overall strength of this effect was also indistinguishable between imagery groups. This suggests that the sensory information represented in early visual cortex was equally important for strong and weak imagers to successfully guide behavior. We thus find no evidence for differences between strong and weak imagers, neither in the encoding of sensory information nor in the degree to which this information is predictive of behavior. These results go against our key prediction from the cognitive-strategies framework of working memory (Pearson & Keogh, 2019), according to which strong imagers should retain higher levels of stimulus information in their early visual cortices during working memory, compared to weak imagers. Our results therefore call into question the importance of experienced imagery vividness in the modulation of early visual cortex recruitment during working memory. Please note that these null effects were based on preregistered analyses and are supported by additional Bayesian analyses.

To our knowledge, this is the first study to specifically investigate the decodability of working memory representations in the context of individual differences in imagery ability. While some studies have considered the relationship between visual imagery and stimulus decoding (Albers et al., 2013; Dijkstra et al., 2017; Dijkstra et al., 2018), they have relied on random samples of participants, potentially not covering the entire spectrum of imagery ability and not addressing the effects of individual differences. One study found that the overlap between imagery and perception signals in early visual cortex is modulated by trial-by-trial imagery measures (Dijkstra et al., 2017). In a later study, the same authors could successfully cross-decode between the neural signatures of weak and strong imagers, indicating that the decodable signal between both groups was similar (Dijkstra et al., 2018). While the second study in particular seems to support our results, caution is advised when comparing results obtained via trial-by-trial measures of imagery with trait measures such as VVIQ scores. Another study has reported a positive relationship between imagery ability and decoding accuracy (Albers et al., 2013), however, note that the authors of that study equated imagery ability with task performance, making this result more analogous to our reported relationship between target reconstruction and behavioral precision. Therefore, our present finding that working memory signals do not seem to depend on imagery vividness is not in direct contradiction to these previous decoding studies.

Importantly, our study was specifically designed to assess the neural encoding of working memory contents, not the neural representations of imagery. If working memory signals in early visual areas were to exclusively reflect imagery, one would predict these working memory signals both to be modulated by imagery ability and to be completely absent for individuals without phenomenal imagery (aphantasics). Our results show that both are not the case. Please note that we are not claiming that visual imagery and visual working memory are never based on the same neural signals. It is possible that imagery in strong imagers recruits the same neural representations that are also used for visual working memory. This would be compatible with findings from previous studies (Albers et al., 2013; Dijkstra. et al., 2017; Dijkstra et al., 2018). However, our finding that even a strong reduction in imagery does not affect the decodable information suggests that these early visual signals are not necessarily tied to imagery. What our data show, is that neural representations of working memory contents are still observable and have a comparable information level even for individuals with weak or absent imagery. Thus, working memory signals can be dissociated from visual imagery in early visual cortex. Note that the current study did not focus on any particular encoding strategy and therefore does not allow any claims about the neural encoding of imagery contents. While it would be interesting to investigate how the strength of imagery representations varies with the vividness of subjectively experienced imagery, this is a question for future research and was not the aim of this study.

Our finding of a close link between sensory information in the delay period and behavioral working memory performance is in line with several previous studies (Bettencourt & Xu, 2016; Ester et al., 2013; Hallenbeck et al., 2021; Harrison & Tong, 2009; Iamshchinina et al., 2021). Based on our highly sensitive method for reconstructing continuous stimulus features from voxel patterns, the neural information explained more than half of the between-subject variance in behavioral performance (see Methods for more details), which further corroborates the link between information encoded in early visual cortex and memorization of visual information across brief delays. Additionally, we found that sensory information was retained only for the cued and thus task-relevant stimulus but was not present for the uncued image. These results are in line with sensory recruitment accounts of working memory (D’Esposito & Postle, 2015), or more generally with a multi-level representation of sensory information across delays (Christophel et al., 2017), according to which cortical areas that are used for the encoding of task-relevant sensory information are also recruited for the brief memorization of that information. This task-dependent retention of information in early visual cortex could point towards some form of *active maintenance* throughout the delay after offset of the stimulus. This could be achieved by neural mechanisms such as recurrent processing within early visual cortex (Lamme & Roelfsema, 2000) or by feedback from higher regions (Gazzaley & Nobre, 2012) and could include short-term synaptic plasticity (Mongillo et al., 2008; Rose et al., 2016). Please note that sensory recruitment does not make any assumptions about the strategy with which sensory information is encoded, i.e., whether it is accompanied by imagery or not.

It is worth pointing out that there has been some debate about the importance of early visual cortex for the generation and maintenance of visual imagery in general. For instance, results from activation-based studies have suggested that imagery effects in early visual cortex might be linked to sensory memory retrieval (Kaas et al., 2010). Further, it has been shown that vivid phenomenal imagery can be preserved in cortically blind patients after strokes to occipital areas (Bartolomeo et al., 1998; Chatterjee & Southwood, 1995; de Gelder et al., 2015), indicating that early visual cortex is not essential for visual imagery. Similarly, lesions in temporal regions have been reported to selectively affect visual imagery but leave visual perception largely preserved (Moro et al., 2008; Thorudottir et al., 2020), which has been taken as evidence that visual imagery depends on a temporal network (Spagna et al., 2021). Together, this would suggest a functional dissociation of early visual cortex and visual imagery (Bartolomeo et al., 2020), with imagery relying on higher-level representations beyond early visual cortex (Bartolomeo, 2008). As a consequence, orientation-specific signals could be maintained in early visual cortex, but weak imagers might not be able to access them to produce phenomenal imagery. On this basis, one could speculate that the weak imagers in our case might have had a deficit in a (potentially temporal) imagery network, whereas working memory performance is based on sensory information that is largely intact. Early visual information would thus be available to solve the working memory task but would not necessarily lead to the experience of imagery. Importantly, however, this is at odds with a large body of behavioral, neuroimaging and brain-stimulation work which suggests a close link between signals in early visual areas and imagery (Albers et al., 2013; Dijkstra et al., 2017; Keogh et al., 2020; Pearson, 2019), a discrepancy which will have to be resolved by future research. Another explanation for our results might be that our participants simply did not use visual strategies at all, or just to a small extent. This would be in direct opposition of the cognitive-strategies framework, which assumes a close correspondence between individual imagery ability and the cognitive strategy used to solve a working memory task (Pearson & Keogh, 2019). Strong imagers usually report to use visual strategies (Bainbridge et al., 2021; Keogh et al., 2021; Logie et al., 2011), and the spontaneous use of visual vs. non-visual strategies by strong and weak imagers has also been confirmed behaviorally, by showing that only strong imagers were affected by distracting visual input during a working memory delay (Keogh & Pearson, 2014). It is therefore unlikely that the strong imagery group in this study relied predominantly on non-visual strategies to solve the task.

One reason for some of the discrepancies in the imagery literature may lie in the different ways in which imagery vividness is quantified across studies (Pearson, 2020). To date, various approaches have been suggested, including self-report questionnaires, trial-by-trial vividness measures (Dijkstra et al., 2017; Dijkstra et al., 2018; Dijkstra et al., 2017) and several measures that are related to certain spontaneous perceptual (Pearson et al., 2008) or physiological (Kay et al., 2022) reactions or anatomical features (Bergmann et al., 2016). It is not yet clear, however, which of these measures provides the best approximation for general individual imagery ability. Some of the more objective measures in particular have been used very rarely and still await calibration with respect to more conventional measures of visual imagery. In contrast, the VVIQ provides a well-established, reliable assessment for individual differences in imagery vividness (Dijkstra et al., 2018; Pearson et al., 2011). VVIQ scores have been shown to successfully capture the relationship between imagery vividness and neural signals (Amedi et al., 2005; Cui et al., 2007; Lee et al., 2012), and people are generally able to provide good metacognitive judgments about their own imagery abilities (Pearson et al., 2011; Rademaker & Pearson, 2012). Further, the VVIQ is closely related to a perceptual priming based measure of imagery ability (Pearson et al., 2008, 2011). In combination with pre-selection and high test-retest reliability, the VVIQ scores should therefore provide a reasonably good estimate of general imagery ability in the two groups recruited for this study.

It is worth mentioning that our reconstruction results might be explained by other factors than orientation-specific visual representations. Please note that in decoding studies it is generally not possible to fully guarantee that information pertains to the features intended by the researcher instead of other latent confounding variables such as spatial attention or motor preparation that co-vary with these features, as we have pointed out previously (Christophel et al., 2017). For example, the distribution of spatial attention can be very different across seemingly homogenous stimulus sets (Liu, 2016; Yun et al., 2013). Thus, when decoding between two object images one might be decoding the spatial distribution of attention rather than the object identity. This could also be the case for the orientation stimuli used here. However, the role of early visual cortex in encoding of orientations as here has long been established both at a cellular level (Hubel & Wiesel, 1968) as well as the population level (Haynes & Rees, 2005; Kamitani & Tong, 2005; Ts’o et al., 1990). Orientation stimuli as here have been used in many cornerstone studies of working memory (Albers et al., 2013; Bae & Luck, 2019; Harrison & Tong, 2009) and imagery (Keogh & Pearson, 2011, 2014; Pearson et al., 2008). Nonetheless, future studies will be needed to test whether all these findings of orientation encoding in early visual cortex during working memory generalize to other stimulus sets.

In conclusion, we show that the active maintenance of stimulus-related information in early visual areas was present also in participants who report a near-absence of visual imagery. The encoding of sensory information and its link to performance was strong and indistinguishable across different levels of imagery. This provides further evidence for the view that the recruitment of early visual cortex for working memory can be dissociated from visual imagery, at least for participants with weak or absent imagery. Thus, informative working memory representations in visual cortex are maintained irrespective of whether a person is able to engage in vivid imagery or not.

## Acknowledgements

Funded by the Deutsche Forschungsgemeinschaft (DFG) Research Training Group 2386 (S.W.); EXC 2002/1 “Science of Intelligence” (S.W., K.G.); SFB 940 “Volition and Cognitive Control” (J.-D.H.); SFB-TRR 295 “Retuning dynamic motor network disorders using neuromodulation” (J.-D.H.); and supported by BMBF and Max Planck Society.

## Author contributions

Conceptualization, S.W., and J.-D.H.; methodology, S.W., T.C., K.G., J.S., and J.-D.H.; investigation, S.W., T.C.; formal analysis, S.W., K.G., and J.-D.H.; software, S.W., K.G., and J.S.; visualization, S.W.; writing, S.W., and J.-D.H.; supervision, J.-D.H.

## Methods

### Data and code availability

Original code, summary statistics describing the reported data and processed datasets which can be used to recreate the figures in this manuscript have been deposited and are publicly available at https://github.com/simonweber91/WM_VI_EVC. Any additional data and information required to reanalyze the data reported in this paper are available from the lead contact upon reasonable request.

### Preregistration

The main analysis workflow of this study (including custom preprocessing steps, parameter choices, ROIs and newly implemented statistical models) was preregistered at https://osf.io/34y9z. The preregistration was submitted after data acquisition, but prior to data processing and analysis. All preregistered analysis procedures were developed and/or optimized on a separate fMRI dataset from a related study (Barbieri et al., 2023). Please note that we did not change any of the preregistered workflows. However, we did perform additional analyses and performed more extended statistical testing (e.g., Bayesian and permutation-based tests) whenever it proved necessary to the quality of the study. All of these additional analyses are indicated as E.A. (“extended analysis”) in this text.

### Recruitment

Two groups of study participants were preselected for the study using an online version of the Vividness of Visual Imagery Questionnaire (VVIQ; Marks, 1973) The questionnaire was implemented and hosted on the online survey platform SoSci Survey (www.soscisurvey.de) and local respondents were recruited via in-house mailing lists for experimental studies, study participant databases and Facebook. Respondents gave informed consent prior to being directed to the questionnaire and again before providing an email address for recruitment at the end of the questionnaire.

We received a total of 263 online responses, 210 of which fulfilled the physiological, medical and demographic criteria for participation in the MRI study. Respondents whose VVIQ scores fell either into the upper or lower quartiles of the response distribution were assigned to the strong and weak imagery groups, respectively, and contacted for participation in the fMRI experiment (Figure 1B). From these groups we recruited a total of 42 fMRI participants. All participants were healthy, right-handed individuals between 18 and 40 years old with no history of neurological or psychiatric disorders. One participant dropped out of the study before completing all scanning sessions. The data of a second participant had to be discarded due to technical issues with the MRI scanner. Therefore, we collected complete datasets of 40 participants (female: 23, age: 28.05 ± 6.064 years), 20 each per experimental group (average VVIQ score; weak: 40.75 ± 11.571; strong: 70.7 ± 3.262).

Participants gave written informed consent prior to the fMRI experiment. They received monetary compensation of 10€/h for the fMRI sessions and a bonus of 10€ for completion of both scanning sessions. Following April 19, 2021, participants were required to present a negative SARS-CoV-19 rapid test result (not older than 24 hours) before entering the MRI facility. To compensate for the additional effort, we paid an additional 20€ for each SARS-CoV-19 rapid test. The study was approved by the ethics committee of the Humboldt-Universität zu Berlin and conducted according to the principles of the Declaration of Helsinki (World Medical Association, 2013).

### Stimuli

The experiment was implemented using MATLAB R2018b (The MathWorks, Inc.) and Psychtoolbox 3 (Brainard, 1997; Kleiner et al., 2007). All stimuli were presented on black background, to avoid residual luminance interfering with potential visual imagery during the delay period (Keogh & Pearson, 2014). For stimulation, we used circular high contrast sine-wave Gabor patches with phase 0, contrast 0.8 and a spatial frequency of 0.02 cycles per pixel. Stimuli were presented inside a circular aperture with an inner diameter of 0.71 dva and an outer diameter of 8.47 dva. A white fixation dot of 0.18 dva was placed at the center of the inner aperture (Figure 1A).

The set of target orientations comprised 40 discrete, equally spaced orientations separated by 180°/40 = 4.5°. To avoid the exact cardinal directions (0°, 45°, 90°, 135°), the orientations were slightly shifted by 1.125°, resulting in a set of orientations between 1.125° and 176.625°. Another set of 40 gratings, which served as distractors, was created by shifting the target orientations by 4.5°/2 = 2.25°, yielding orientation stimuli between 3.375° and 178.875°. This ensured that (i) target and distractor orientations were never exactly the same and (ii) both sets of orientations avoided the exact cardinal directions. Since we presented 40 trials in each run (see below), each target and distractor orientation was shown once during each run, in randomized order. Accordingly, target and distractor orientations were counterbalanced across runs. The starting orientation of the probe grating was randomly selected from a uniform distribution between 0° and 180° on each trial.

To avoid afterimages, we used a custom dynamic noise mask (Figure 1A). For each presentation of the mask, we initialized a 42-by-42 array of an equal number of black and white squares. Each time the screen was refreshed (refresh rate: 60 Hz), the array was scrambled along the rows and columns and smoothed by convolving it with a 2 x 2 box blur kernel. This created a highly dynamic noise mask that reliably suppressed afterimages of the high-contrast gratings. Masks were presented inside the same circular aperture as the stimuli.

### fMRI task

The visual stimuli were presented on an MRI-compatible monitor (dimensions: 52 x 39 cm, resolution: 1024 x 768 px), positioned at the far end of the scanner bore, and viewed via an eye-tracking compatible mirror mounted on top of the head-coil. The distance between the eyes and the center of the monitor was 158 cm.

Each trial of the experiment started with the presentation of a central fixation dot which remained visible throughout the entire trial (Figure 1A). Participants were instructed to fixate the dot at all times. After 0.4 s, participants were sequentially presented with two gratings (see above), one serving as the target and the other as the distractor. Each grating was shown for 0.4 s, followed by 0.4 s of a dynamic, high-contrast noise mask to avoid after-images. After the second mask a numerical retro-cue (0.4 s) was presented at the location of the fixation dot, indicating to the participants to remember the orientation of either the first (“1”) or second (“2”) grating during the subsequent delay period. The delay period lasted for 10 s, during which only the fixation dot remained visible on the screen. After the delay, a probe grating with random starting orientation appeared for 3.2 s. Participants were asked to adjust the orientation of the probe grating in a way that it corresponded to the remembered (target) orientation, using two buttons with the index and middle fingers of their right hand. After adjustment, participants had to confirm their response by pressing a button with the index finger of their left hand. If the response was completed within the time-window of 3.2 s, the fixation dot turned green for the remainder of the response period as visual feedback. If participants failed to provide a response in time, a small “X” was presented at the location of the fixation dot for 0.4 s. Trials were separated by a variable inter-trial interval (ITI) of 3.6 ± 1.6 s. Participants completed 40 trials per run and a total of 8 runs, equally split across 2 fMRI sessions on separate days, resulting in 320 trials per participant.

### MRI data acquisition

MRI data were collected with a 3-Tesla Siemens Prisma MRI scanner (Siemens, Erlangen, Germany) using a 64-channel head coil. At the beginning of each session, we recorded a high-resolution T1-weighted MPRAGE structural image (208 sagittal slices, TR = 2400 ms, TE = 2.22 ms, TI = 1000 ms, flip angle = 8°, voxel size = 0.8 mm^2^ isotropic, FOV = 256 mm). On each of the two days, this was followed by four experimental runs, for each of which we recorded a series of 965 T2-weighted functional images using a multi-band accelerated EPI sequence with a multiband factor of 8 (TR = 800 ms, TE = 37 ms, flip angle = 52°, voxel size = 2 mm^2^ isotropic, 72 slices, 1.9 mm inter-slice gap), resulting in a duration of 12:52 min per run. The first four TR of each sequence were discarded.

### Eye-tracking

We used an EyeLink 1000 Plus (SR-Research) eye-tracker to record gaze position and pupil size of the dominant eye of each participant during the experimental runs. The tracker was positioned at the far end of the scanner bore (eye-lens-distance: 85 cm) on a long-distance mount and was calibrated once at the beginning of each session. Due to technical difficulties, we were only able to record eye-tracking data of 26 participants (13 per experimental group).

### Post-experiment questionnaires

After the second session, participants completed three questionnaires: (i) the Vividness of Visual Imagery Questionnaire (VVIQ, as a post-experimental reference); (ii) the Object-Spatial Imagery Questionnaire (OSIQ; Blajenkova et al., 2006) a 30-item questionnaire probing the strength of visual and spatial imagery; and (iii) a purely heuristic strategy questionnaire, asking (on a 5-point scale) for the degree to which they had used specific mnemonic strategies to remember the target orientations and complete the task, including visual, verbal, spatial, reference to cardinal directions, reference to a clock face, some kind of individual code, or other.

### Behavioral data analysis

Behavioral responses were modeled by fitting a von Mises mixture model (vMMM) to the distribution of behavioral response errors (see Töpfer et al., 2022; original code available at https://github.com/JoramSoch/RDK_vMMM). The model is inspired by previous work on modelling detections from working memory with similarly continuous features (Zhang & Luck, 2008). In our case, we assume that on every trial participants either *detect* the target (responses to target orientations, assumed to follow a von Mises distribution with mean 0° plus bias µ and precision *κ*), make a *swap error* (responses to distractor orientations, following the same assumptions as detections) or *guess* (assumed to follow a continuous uniform distribution between −90° and +90°). Each of these three potential trial-wise outcomes (detections, swaps and guesses) has an associated probability distribution indicating how probable each potential response angle is, given the orientation of the stimulus (i.e., target and distractor). The overall response distribution is considered a linear combination of these three individual event probability distributions with associated probabilities as mixture coefficients *r*_1_, *r*_2_ and *r*_3_.

According to this approach, the probability of observing a specific response evaluates to

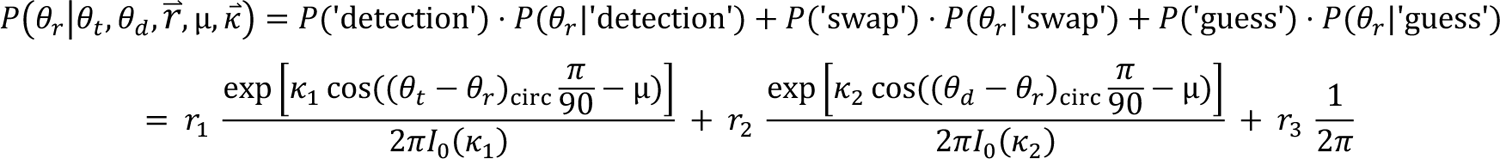

where θ_r_ is the reported orientation in degrees; θ_t_, θ_d_ and are the target and distractor orientations in degrees, respectively; *r*^→^ is a vector containing *r*_1_, *r*_2_and *r*_3_, the event probabilities for the three model components (detections, swap errors and guesses); *K*^→^ is a vector containing *K*_1_ and *K*_2_, the precisions for detections and swap errors, respectively; µ is the response bias and *I*_’_(*K*_i_) is the modified Bessel function of order 0. As *K*_1_ reflects the width of the response distribution for target detections, we report this parameter as our key measure for behavioral precision.

### fMRI preprocessing

Processing and analysis of fMRI data was performed in MATLAB 2021b, using SPM12, The Decoding Toolbox (Hebart et al., 2015) and custom scripts (see below). MR images were converted into NIfTI format for further processing. Before the analysis, BOLD images were spatially realigned and resliced. The T1 image of each session was coregistered to the first image of the respective BOLD series. We then calculated normalization parameters to the Montreal Neurological Institute (MNI) standard space. These were used to project probabilistic maps of our regions of interest (ROIs) into the native space of each individual participant to guide voxel selection during the reconstruction analysis (see below). Following realignment, the time series of each voxel’s raw data were temporally detrended, to remove slow signal drifts that accumulate across a given run. This was implemented using cubic spline interpolation (modifying an existing algorithm; Tanabe et al., 2002). The time series of voxel data for a given run was separated into 40/2 = 20 segments of equal size. The data from each segment was averaged to create query points (nodes), which were then used for cubic spline interpolation, creating a smooth function modeling the slow signal drifts in the voxel data across the run. The number of nodes was specifically set to half the number of trials per run, to avoid the modeling (and thereby, removal) of within-trial effects. The drift-estimate was then subtracted from the voxel data. This procedure was repeated for every voxel and every run. After detrending, we applied temporal smoothing to the data by running a moving average of width 3 TR across the data of each run.

To increase the signal-to-noise ratio for samples from trials with neighboring stimulus orientations, we developed a method that we refer to as “feature-space smoothing”. Feature-space smoothing accounts for the assumption that, in a feature-continuous stimulus space, samples that lie closely together in feature space (e.g., neighboring orientations) should produce a similar neural response and therefore a similar voxel signal. By reducing the contribution of noise to the measurements of neighboring samples, it should be possible to increase the amount of information represented in the voxel signal across the feature space. We addressed this issue by using a gaussian smoothing kernel to compute a weighted average of the voxel signal corresponding to a given orientation and its neighbors (Figure S3). This means that samples close to a given orientation in feature space contribute more to the resulting average than those further away. The number (or distance) of samples included in the average is determined by the width (full width at half maximum, FWHM) of the smoothing kernel. Please note that we confirmed through simulations that feature-space smoothing can substantially increase signal-to-noise ratio and thereby reconstruction accuracies without producing spurious above-chance accuracies in the case of null data (Figure S3). In this study, we used nested cross-validation across subjects to determine the optimal kernel width for each participant (see below). Please note that all these approaches for temporal detrending and feature-space smoothing were developed and optimized on a separate dataset (from a related study; Barbieri et al., 2023) and both were pre-registered and checked for artifacts or spurious effects.

### Early visual cortex ROI

As our goal was to determine the strength of working memory representations in visual sensory stores depending on visual imagery vividness, we restricted our analysis to visually driven voxels in early visual cortex (V1, V2, V3). These regions have been shown repeatedly to similarly encode working memory representations of orientation (and other visual) stimuli (Christophel et al., 2012; Christophel & Haynes, 2014; Harrison & Tong, 2009; Serences et al., 2009). In a first step we combined the probabilistic anatomical maps of V1, V2 and V3 (Wang et al., 2015) to create a combined map in standard space, collapsing across left and right hemispheres. We then transformed this map into the native space of each participant, by applying the inverse normalization parameters estimated during preprocessing. The individual maps were then thresholded at 0.1, to exclude voxels that had a less than 10 % probability of being part of a given area, and binarized. This resulted in an average ROI size of 5938.6 ± 858.45 voxels. In a second step we identified visually driven voxels within that ROI. For this, we estimated a GLM with regressors for all trial events (target, distractor, cue, delay and probe, plus 6 head motion realignment parameters as regressors of no interest). Regressors were convolved with a canonical hemodynamic response function. We then calculated a contrast for the target regressor (vs. an implicit baseline), in order to determine voxels with significant activation in response to the target, irrespective of orientation. The resulting statistical parametric maps were then used in combination with the individual anatomical ROIs for voxel selection in the multivariate reconstruction analysis. For this, we selected the voxels rank-ordered by their respective *t*-score (from the unspecific target contrast) within the anatomical ROI for each individual. The cutoff yielding the exact number of voxels used for reconstruction was determined via nested cross-validation across subjects (see below).

### Orientation reconstruction from fMRI data

The aim of our reconstruction analysis was to predict the angle of the orientation stimulus from the multivariate signal of the preprocessed raw data in the early visual cortex ROI. Note that the space of orientations is circular between 0° and 180°. To account for this, we implemented periodic support vector regression (pSVR), a periodic extension of the SVR (Drucker et al., 1996). First, we projected the angular labels into a periodic space by calculating two sinusoids in the range [0°, 180°). Both functions had an amplitude of 1 and a period of 180°, so that one period spanned the entire label space. One function was shifted by 45°, so that the combination of both periodic functions coded for the linear label scale (Figure S4). This is the 180°-equivalent to the way sine and cosine functions between 0° and 360° code for the angles on a unit circle.

Next, we individually predicted each set of labels from the multivariate voxel pattern using the LIBSVM (Chang & Lin, 2011) implementation of SVR with a non-linear radial basis function (RBF) kernel, via a leave-one-run-out cross-validation. Before prediction, the voxel signals in the training data were rescaled to the range [0, 1]. The scaling parameters were then applied to the test data (“across-scaling”; Hebart et al., 2015).

After prediction of both sets of periodic labels (*x*_i_, *y*_i_) we computed the reconstructed angular orientation θ_i_ using the four-quadrant inverse tangent:

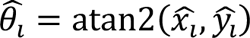

The analysis was repeated for the 30 TRs (24 s) following delay-onset, for each TR individually. This allowed for a time-resolved estimation of how orientations were represented in the visual cortex across the entire trial.

### Reconstruction performance evaluation

To evaluate the accuracy of the orientation reconstruction, we computed the feature-continuous accuracy (FCA). FCA is a rescaling of the absolute angular deviation (between predicted and true label) into the range 0-100 % and can be calculated, for the case of stimuli that are 180°-periodic, as (Pilly & Seitz, 2009)

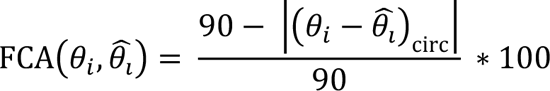

where θ_i_ is the true orientation in the *i*-th trial and θ_i_ is the associated reconstructed orientation. This trial-wise measure of reconstruction performance can be easily interpreted as a feature-continuous analogue to the accuracy measure of more conventional classification approaches: a value of 100 % means that there is no deviation between true and reconstructed orientations, i.e., perfect reconstruction; 50 % means deviation of 45°, which for circular orientation data is equivalent to guessing and can be considered as the chance-level; and 0 % means that reconstructed and true orientations are exactly orthogonal. FCA can be averaged to quantify reconstruction accuracy across trials.

For behavioral responses, the orientation labels may not be uniformly distributed across the orientation space, but clustered around, for example, cardinal axes. In a reconstruction setting, this would be analogous to a classification case with unequal (or unbalanced) numbers of classes, where the predictive model can exploit the uneven distribution of classes to simply predict the more frequent class more often. To account for this potential source of bias, we calculated a balanced FCA (BFCA). BFCA is an extension of the concept of balanced accuracy (Brodersen et al., 2010) for continuous variables. It is calculated by computing the integral of the trial-wise FCA from 0° to 180° (i.e., the orientation-space), using trapezoidal numerical integration across the sorted true and reconstructed orientations: (Barbieri et al., 2023)

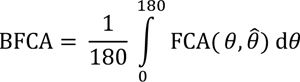

The process of integration assigns lower weights to the FCA values in the well-populated parts of the label-distribution and higher weights to the less populated parts. Thus, BFCA is a non-trial-wise measure of reconstruction performance, which accounts for the potential bias in FCA caused by non-uniformly distributed labels. We report BFCA as our key measure for reconstruction accuracy. Note that this approach has been previously tested to exclude the possibility of artifactual results.

### Parameter optimization

As mentioned above, we used an across-subjects nested cross-validation to determine the optimal values of two parameters for each participant individually: (i) the width of the gaussian kernel used for feature-space smoothing, and (ii) the number of voxels entered into the analysis. For (i), we chose FWHM values between 0° (i.e., no smoothing) and 90°, in steps of 10°. Thus, we had a set of 10 possible kernel widths for smoothing. For (ii), we chose voxel counts between 250 and 2500, in steps of 250. This resulted in a set of 10 possible voxel counts. To select the specific voxels entered into the analysis, we first masked the individual target-versus-baseline *t*-maps with the warped anatomical ROIs (see above) and then selected the n voxels with the highest *t*-scores within those ROIs, with n representing a number from the set of possible voxel counts. The reconstruction analysis was then run for every combination of FWHM values and voxel counts.

After reconstruction, we determined the optimal parameters for each subject in the following way: First, we calculated the mean BFCA across all remaining subjects for every parameter combination, resulting in one value per combination and time point. Second, we averaged across the preregistered delay-period TRs (TRs 6-15 following delay onset), as we were specifically interested in potential group differences during this time window. This yielded one BFCA value per parameter combination, specifically for the entire delay period. The parameter combination that yielded the highest BFCA was then assigned to the left-out subject. This was repeated for every subject and resulted in an average FWHM value of 74.5 ± 9.04 and an average voxel count of 1750 ± 211.83.

### Statistical Testing

As we were specifically interested in potential group differences during the delay-period, statistical testing for differences between the strong and weak imagery groups was based on the time points in the trial which most likely only reflect delay period activity. Since the canonical hemodynamic response has a buildup of ∼5 seconds, we considered the TRs 6-15 in the 30 TR timeframe that we analyzed, corresponding to a time window of 4 s after delay onset to 2 s after probe onset (please note that this time window is 0.4 s shorter than described in the preregistration, as the preregistered time window would have resulted in 10.5 instead of 10 TRs). This preregistered time window should avoid the leaking of stimulus- or probe-representations into the delay-period analysis.

We used two-tailed two-sample *t*-tests to test for potential differences in the reconstruction scores between the experimental groups. Further, we calculated Pearson’s *r* to assess the correlation between outcome variables (E.A.).

*Cluster-based permutation approach (E.A.)* We were interested at which time points during the trial we could detect significant above-chance reconstruction accuracy. To account for the multiple-comparisons (Groppe et al., 2011) and autocorrelation (Purdon & Weisskoff, 1998) issues that arise for such time-resolved analyses, we adopted a non-parametric cluster-based permutation approach (Bullmore et al., 1999; Groppe et al., 2011; Maris & Oostenveld, 2007). This procedure was performed after the parameter optimization described above, to restrict the time-consuming permutation analysis to one set of parameters per subject. We repeated this approach separately for each reconstructed label type: target, distractor, probe and reported orientation.

*Bayesian tests (E.A.).* As our results indicated no significant differences between our two groups, we used Bayesian hypothesis tests to assess the evidence for this absence. Bayesian hypothesis tests are used to describe the probability of observing the measured data under the null and alternative hypothesis, respectively (Keysers et al., 2020). This likelihood is quantified using the Bayes factor (BF), a continuous measure of evidence for either hypothesis. Specifically, we used two Bayesian hypothesis tests to assess the evidence for absence of effects: First, in the case of non-significant group-comparisons, we performed follow-up Bayesian independent *t*-tests, using a Cauchy distribution with scale parameter *r* = 0.707 as the prior distribution (Morey & Rouder, 2011). Second, in the case of non-significant correlations, we performed Bayesian correlation with a stretched beta prior of width *κ* = 1. All Bayesian hypothesis tests were performed in the open-source software JASP (Love et al., 2019).

### Orientation reconstruction from eye-tracking data

Participants were instructed to maintain fixation at all times during the experiment. It is at least theoretically conceivable that participants might have used an eye-movement-based strategy to remember target orientations. Eye-movements have also been shown to modulate visual responses in the brain (Merriam et al., 2013). To account for these potentially confounding factors, we investigated whether the gaze position across the trial held information about the target orientation. For this, we subjected the recorded x and y ordinates of 26 participants (for which complete sets of eye-tracking data were available) to the same reconstruction analysis as the fMRI data.

Preprocessing of eye-tracking data was performed in MATLAB using functions from the Fieldtrip toolbox (Oostenveld et al., 2011), code adapted from prior work (Urai et al., 2017) and in-house code. Blinks were linearly interpolated and bandpass filtered between 5 Hz (high-pass) and 100 Hz (low-pass). For each trial, we extracted 15 s worth of data following the onset of the first grating. The data from each run was detrended using the same cubic spline interpolation as described above (see Preprocessing of fMRI data). We then downsampled the data by a factor of 10, resulting in 1500 time points per trial.

After preprocessing, we entered the data into the same pSVR reconstruction analysis as the fMRI data, using the x and y ordinates of the gaze position as input instead of voxel signal, and evaluated the reconstruction by calculating the BFCA. As with the fMRI data, we tested for clusters of above-chance time points using the cluster-based *t*-mass permutation approach described above.

### Feature-space smoothing simulation

To demonstrate how feature-space smoothing can increase signal-to-noise ratio (SNR) and increase accuracy in a continuous reconstruction setting, we simulated fMRI data with varying amounts of SNR and used different levels of feature-space smoothing before reconstruction. Following the specifics of our experiment, we simulated data comprising 8 runs with 40 trials each, for 250 voxels. The measured response of voxel *i* in trial *j* was generated as

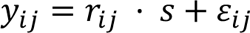

where *r*_ij_ is the actual response of voxel *i* in response to the orientation shown in trial *j*, *s* is a scaling factor controlling the ratio of signal and noise, and ε_ij_ is sampled from a standard normal distribution.

To simulate the voxel responses, we assumed a population of idealized voxels, where each voxel would exhibit a distinct periodic tuning profile in response to angular orientation. The tuning profile *Z*_i_ for each voxel *i* was sampled from a multivariate normal distribution

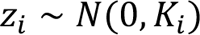

where *K*_i_ specifies the voxels’ periodic covariance kernel. This kernel *K*_i_ is given by

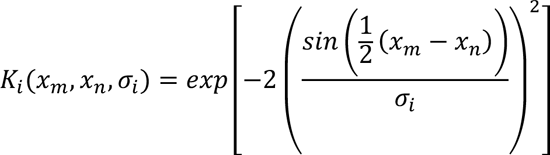

where *x* is a *p* × 1 vector specifying a grid of possible orientations, such that *x*_m_, *x*_n_ ∈ [0,2π), *p* is controlling the number of unique, equally spaced values from the feature space; and σ_i_ is the voxel’s unique tuning function smoothness parameter. For this simulation, the smoothness of each voxel was sampled from a gamma distribution: σ_i_ ∼ Γ(2,2).

Thus, voxel- and trial-wise responses could be sampled as

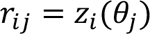

where *x*_i_ is the orientation presented during the *j*-th trial and orientation labels were drawn from a uniform distribution: θ_i_ ∼ *U*(0,2π).

For the SNR-controlling factor *S*, we chose 10 values between 0.1 and 1, equally spaced by 0.1, as well as 0 (i.e., pure noise). Before reconstruction, we used feature-space smoothing on the data, for FWHM values between 0° (i.e., no smoothing) and 360°, equally spaced by 10°. This resulted in 11 SNR levels and 37 smoothing levels. After pSVR reconstruction, we calculated BFCA as our measure of accuracy. The simulation was repeated 1000 times for each parameter combination. The results of this simulation are summarized in Figure S3.

## Supplemental Information

**Figure S1.**
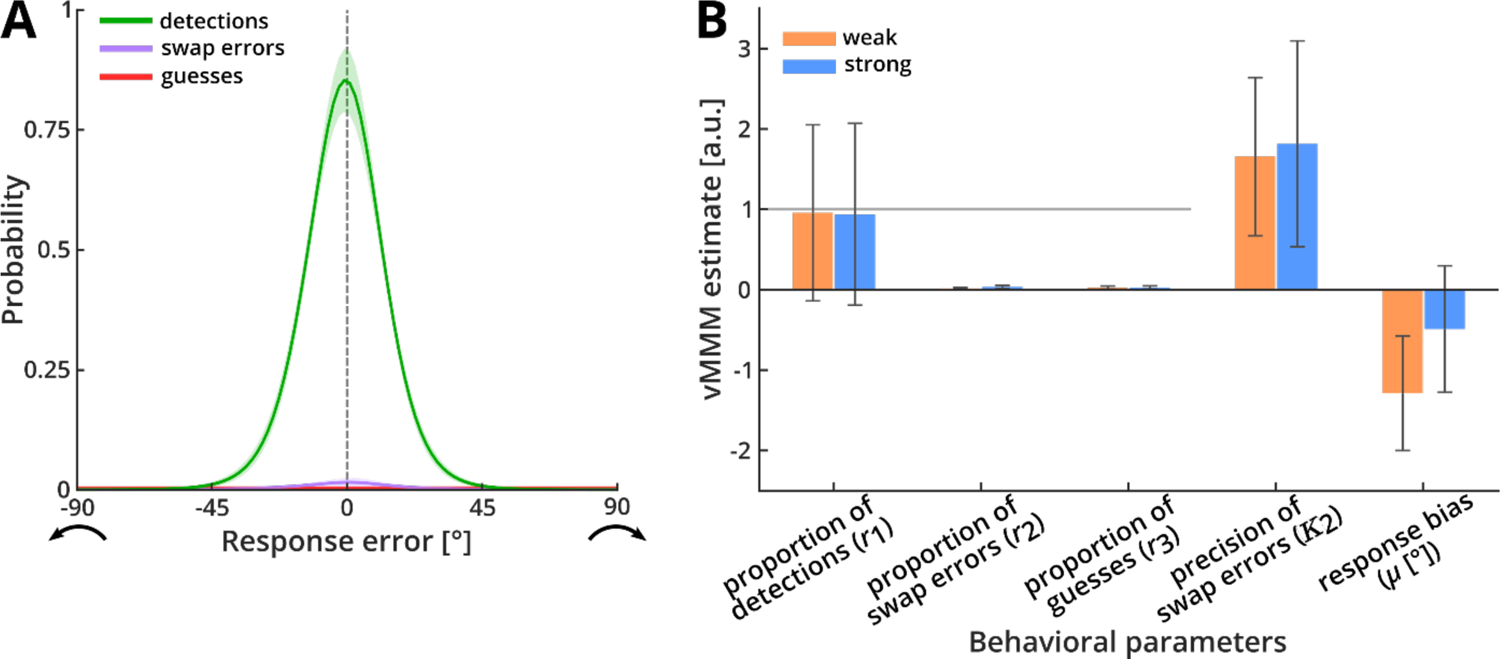
Von Mises mixture model (vMMM) fit of behavioral responses. **(A)** The distribution of behavioral responses was modeled as a combination of the three model components: detections (responses to target orientations, assumed to follow a von Mises distribution with mean 0° plus bias µ and precision *κ*; green), swap errors (responses to distractor orientations, following the same assumptions as detections; purple) and guesses (assumed to follow a continuous uniform distribution between −90° and +90°; red). These components were weighted by individual event probabilities (mixture coefficients) *r*_1_, *r*_2_and *r*_3_, respectively. Participants correctly responded to the target direction in 94.7 % of trials (*r*_1_ = 0.947 ± 0.063), and only infrequently made swap errors (*r*_2_ = 0.026 ± 0.034) or guesses (*r*_3_ = 0.027 ± 0.041). Responses to targets were precise (*K*_1_ = 5.673 ± 2.377), while responses to the distractor, where present, were imprecise (*K*_i_ = 1.735 ± 2.41). There was a small but significant bias to respond anti-clockwise of the target (*μ* = −0.889 ± 1.635°; *t*_(39)_ = −3.437, *p* = 0.0014, two-tailed; see also Figure 1C). **(B)** Estimated vMMM parameters for strong and weak imagers separately. There was no significant difference between the two groups for any of the estimated parameters (*r*_1_: *t*_(38)_ = −0.925, *p* = 0.361; *r*_2_: *t*_(38)_ = 1.585, *p* = 0.121; *r*_3_: *t*_(38)_ = 0.108, *p* = 0.914; *K*_i:_ *t*_(38)_ = −0.207, *p* = 0.837; *μ*: *t*_(38)_ = 1.574, *p* = 0.124, all two-tailed; see Figure 1D for *K*_!_).

**Figure S2.**
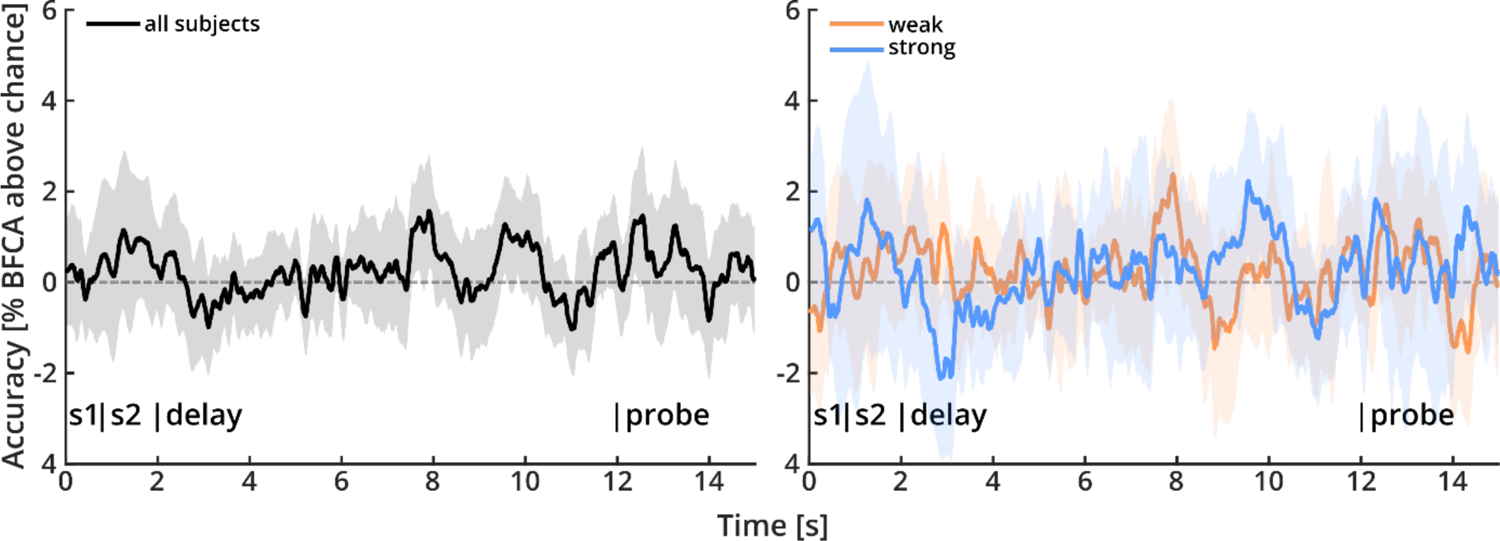
Target reconstruction from eye-tracking data. Reconstruction of target orientation from gaze position across the trial, for all subjects (left panel) and separated by groups (right panel). There were no temporal clusters with significantly above-chance BFCA, suggesting that participants did not systematically use gaze position to maintain target orientation across the delay period. Shaded areas indicate 95 % confidence intervals.

**Figure S3.**
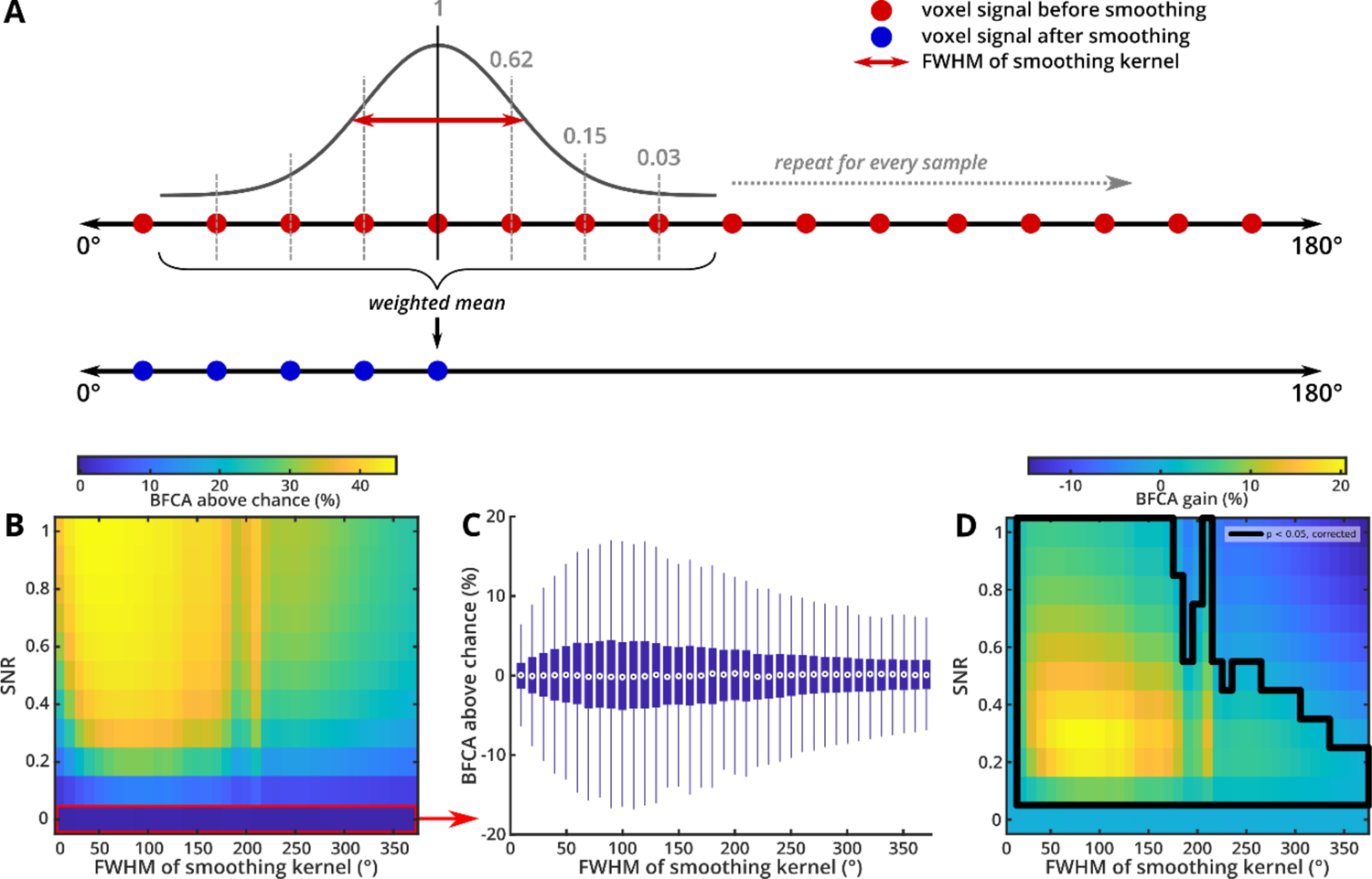
Schematic representation of feature-space smoothing and simulation results. **(A)** We used a Gaussian smoothing kernel to compute a weighted average from the voxel signal of samples lying closely together in feature space. Samples close to a given orientation in feature-space therefore contribute more to the resulting average than those further away. The full width at half maximum (FWHM) of the smoothing kernel controls the smoothing range, i.e., the number (or distance) of samples that are included in the weighted average. We used FWHM values between 0° (no smoothing) and 90° in steps of 10° and determined the optimal kernel width for each participant via nested cross-validation across subjects. Note that this was done (a) at the level of the input data to the analysis, not the results, (b) for training and test data separately, and (c) was confirmed not to produce artifacts or spurious results by extensive simulations (see (C) and Extended Methods). **(B)** We simulated data with varying levels of SNR and used feature-space smoothing with different kernel widths (measured as FWHM in degrees) before reconstruction of the underlying signal. The plot shows BFCA for all parameter combinations, averaged across 1000 repetitions. **(C)** BFCA across smoothing levels, for the pure noise condition. BFCA remained at chance-level across all levels of smoothing (all *p* > 0.25) and BFCA for any smoothing condition did not differ from the no-smoothing condition (all *p* > 0.15). **(D)** BFCA gain compared to no smoothing, averaged across all 1000 repetitions. The first column corresponds to baseline, i.e., zero smoothing. In the signal conditions (SNR > 0), feature-space smoothing was able to reliably increase BFCA compared to no smoothing. The effect was strongest for smoothing kernel widths between 30° and 170°, where we observed increases in accuracy of up to 20 %. Generally, the effect of feature space smoothing was stronger for data with low SNR (orange-yellow area). In cases of extremely high kernel-width and comparatively high SNR (i.e., SNR > 0.6 and FWHM > 220°), feature-space smoothing had a detrimental effect, meaning that BFCA was decreased compared to no smoothing (dark blue area). Please note, however, that kernel-widths this high do not make any sense for real-world applications and were only included for the purpose of demonstration. We conclude that feature-space smoothing is a powerful preprocessing technique to increase SNR in a feature-continuous reconstruction setting. As the optimal kernel-width for smoothing depends on the specific data and SNR, we recommend using nested cross-validation to determine the optimal FWHM value, similar to the approach described in the main text.

**Figure S4.**
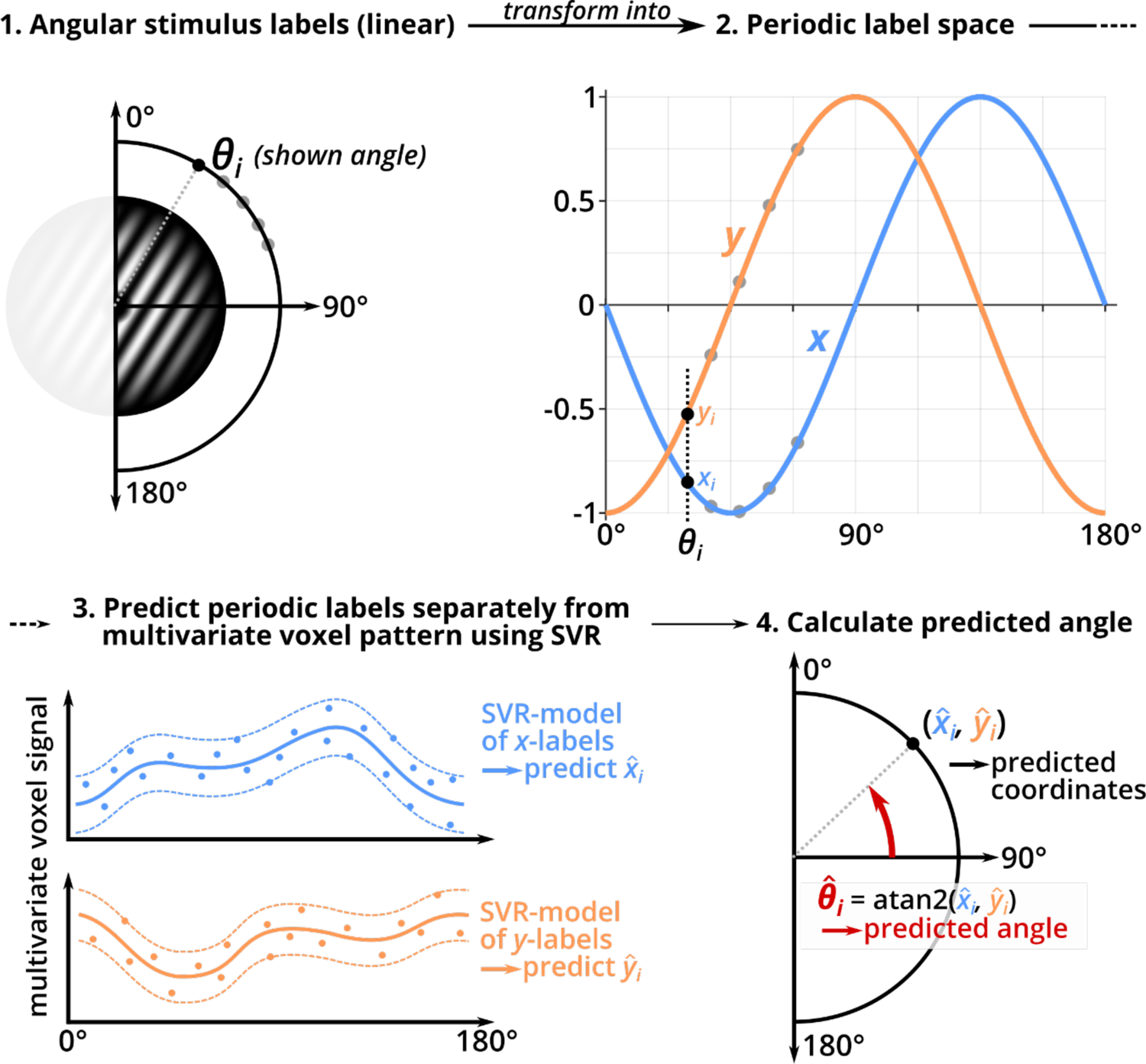
Schematic representation of periodic support vector regression (pSVR). The aim of our reconstruction analysis was to predict an angular label between 0° and 180° from the multivariate voxel signal in response to a stimulus grating with the respective orientation. However, the linear scale of orientation labels (from 0° to 180°) does not reflect the periodic nature of the stimulus (i.e., 0° and 180° are identical). To account for this, we projected the angular labels into a periodic space by fitting two sinusoids into the range [0, 180). Both functions had an amplitude of 1 and a period of 180°, so that one period spanned the entire label space. One function was shifted by 45°, so that the combination of both periodic functions coded for the linear label scale. This is equivalent to the way sine and cosine functions between 0 and 360° code for the angles on a unit circle. We trained and tested a multivariate SVR model for both periodic label sets (*x, y*) separately. From the combination of the predicted periodic labels, we then reconstructed a predicted angular label using the four-quadrant inverse tangent. The predicted orientation was then compared to the true orientation to derive BFCA, our measure of reconstruction accuracy.

---

**Table S1:**
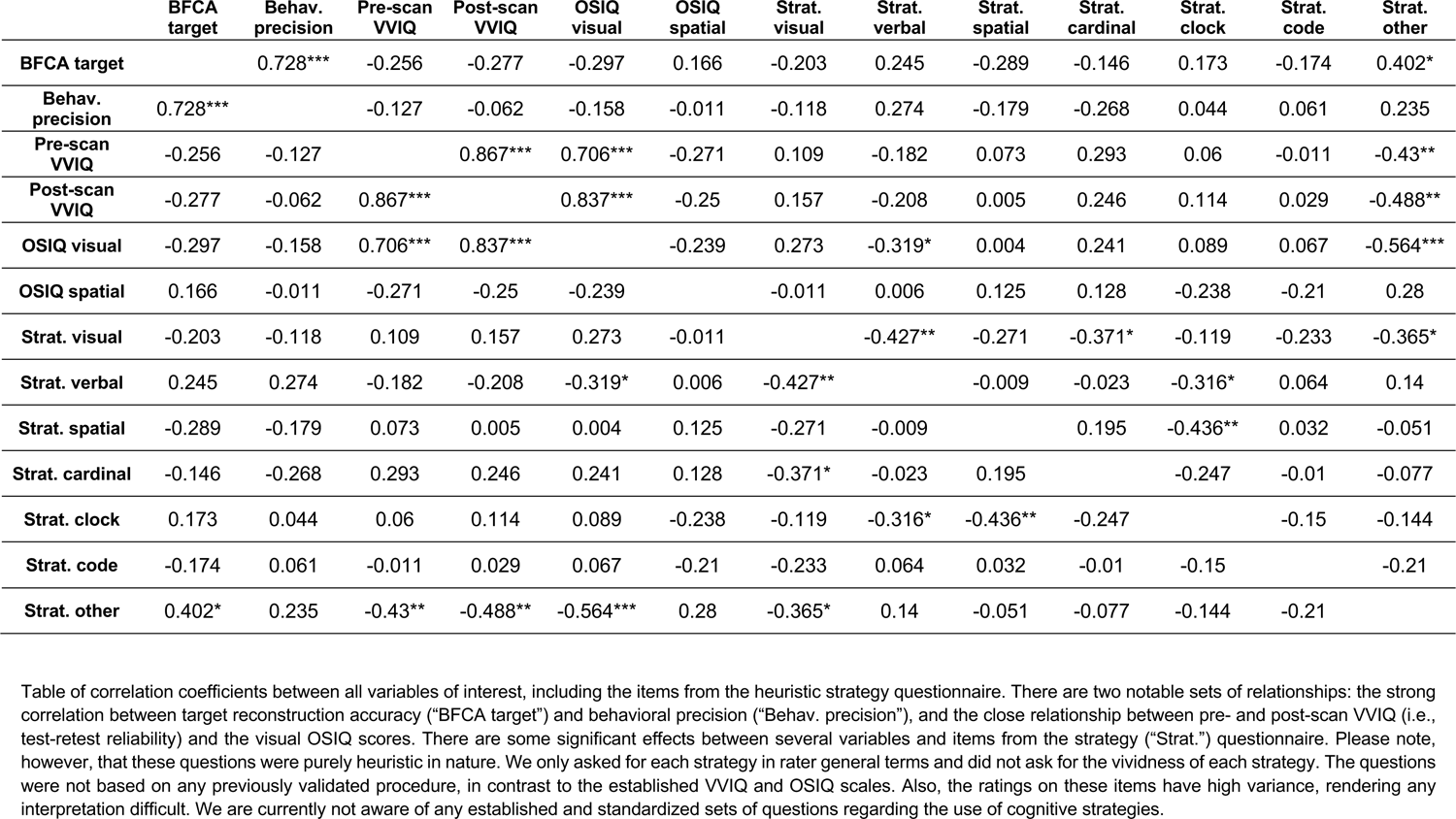
Correlation table of all variables of interest (and strategy questionnaire).

|  | **BFCA target** | **Behav. precision** | **Pre-scan VVIQ** | **Post-scan VVIQ** | **OSIQ visual** | **OSIQ spatial** | **Strat. visual** | **Strat. verbal** | **Strat. spatial** | **Strat. cardinal** | **Strat. clock** | **Strat. code** | **Strat. other** |
| --- | --- | --- | --- | --- | --- | --- | --- | --- | --- | --- | --- | --- | --- |
| **BFCA target** |  | 0.728*** | -0.256 | -0.277 | -0.297 | 0.166 | -0.203 | 0.245 | -0.289 | -0.146 | 0.173 | -0.174 | 0.402* |
| **Behav. precision** | 0.728*** |  | -0.127 | -0.062 | -0.158 | -0.011 | -0.118 | 0.274 | -0.179 | -0.268 | 0.044 | 0.061 | 0.235 |
| **Pre-scan VVIQ** | -0.256 | -0.127 |  | 0.867*** | 0.706*** | -0.271 | 0.109 | -0.182 | 0.073 | 0.293 | 0.06 | -0.011 | -0.43** |
| **Post-scan VVIQ** | -0.277 | -0.062 | 0.867*** |  | 0.837*** | -0.25 | 0.157 | -0.208 | 0.005 | 0.246 | 0.114 | 0.029 | -0.488** |
| **OSIQ visual** | -0.297 | -0.158 | 0.706*** | 0.837*** |  | -0.239 | 0.273 | -0.319* | 0.004 | 0.241 | 0.089 | 0.067 | -0.564*** |
| **OSIQ spatial** | 0.166 | -0.011 | -0.271 | -0.25 | -0.239 |  | -0.011 | 0.006 | 0.125 | 0.128 | -0.238 | -0.21 | 0.28 |
| **Strat. visual** | -0.203 | -0.118 | 0.109 | 0.157 | 0.273 | -0.011 |  | -0.427** | -0.271 | -0.371* | -0.119 | -0.233 | -0.365* |
| **Strat. verbal** | 0.245 | 0.274 | -0.182 | -0.208 | -0.319* | 0.006 | -0.427** |  | -0.009 | -0.023 | -0.316* | 0.064 | 0.14 |
| **Strat. spatial** | -0.289 | -0.179 | 0.073 | 0.005 | 0.004 | 0.125 | -0.271 | -0.009 |  | 0.195 | -0.436** | 0.032 | -0.051 |
| **Strat. cardinal** | -0.146 | -0.268 | 0.293 | 0.246 | 0.241 | 0.128 | -0.371* | -0.023 | 0.195 |  | -0.247 | -0.01 | -0.077 |
| **Strat. clock** | 0.173 | 0.044 | 0.06 | 0.114 | 0.089 | -0.238 | -0.119 | -0.316* | -0.436** | -0.247 |  | -0.15 | -0.144 |
| **Strat. code** | -0.174 | 0.061 | -0.011 | 0.029 | 0.067 | -0.21 | -0.233 | 0.064 | 0.032 | -0.01 | -0.15 |  | -0.21 |
| **Strat. other** | 0.402* | 0.235 | -0.43** | -0.488** | -0.564*** | 0.28 | -0.365* | 0.14 | -0.051 | -0.077 | -0.144 | -0.21 |  |

Table of correlation coefficients between all variables of interest, including the items from the heuristic strategy questionnaire. There are two notable sets of relationships: the strong correlation between target reconstruction accuracy (“BFCA target”) and behavioral precision (“Behav. precision”), and the close relationship between pre- and post-scan VVIQ (i.e., test-retest reliability) and the visual OSIQ scores. There are some significant effects between several variables and items from the strategy (“Strat.”) questionnaire. Please note, however, that these questions were purely heuristic in nature. We only asked for each strategy in rather general terms and did not ask for the vividness of each strategy. The questions were not based on any previously validated procedure, in contrast to the established VVIQ and OSIQ scales. Also, the ratings on these items have high variance, rendering any interpretation difficult. We are currently not aware of any established and standardized sets of questions regarding the use of cognitive strategies.

**Table S2:** Descriptive statistics for all variables of interest (and strategy questionnaire).

|  | **BFCA**<br>**target** | **Behav.**<br>**precision** | **Pre-scan**<br>**VVIQ** | **Post-scan**<br>**VVIQ** | **OSIQ**<br>**visual** | **OSIQ**<br>**spatial** | **Strat.**<br>**visual** | **Strat.**<br>**verbal** | **Strat.**<br>**spatial** | **Strat.**<br>**cardinal** | **Strat.**<br>**clock** | **Strat.**<br>**code** | **Strat.**<br>**other** |
| --- | --- | --- | --- | --- | --- | --- | --- | --- | --- | --- | --- | --- | --- |
| **Mean** | 12.21 | 5.673 | 55.725 | 58.25 | 49.525 | 44.225 | 0.285 | 0.141 | 0.189 | 0.071 | 0.186 | 0.054 | 0.073 |
| **Standard**<br>**deviation** | 9.768 | 2.377 | 17.332 | 15.834 | 12.878 | 8.636 | 0.217 | 0.13 | 0.119 | 0.115 | 0.179 | 0.092 | 0.121 |
| **Skewness** | 0.484 | 1.143 | -0.641 | -1.055 | -1.048 | -0.187 | 1.664 | 0.453 | 0.062 | 1.758 | 0.435 | 1.335 | 1.407 |
| **Kurtosis**<br>**(excess)** | -0.862 | 1.519 | -0.708 | 0.264 | 0.226 | -0.658 | 3.727 | -0.8 | -0.066 | 3.108 | -0.93 | 0.154 | 0.576 |

Table of mean, standard deviation, skewness and excess kurtosis for all variables of interest, including the items from the heuristic strategy questionnaire.

## References

Albers, A. M., Kok, P., Toni, I., Dijkerman, H. C., & de Lange, F. P. (2013). Shared Representations for Working Memory and Mental Imagery in Early Visual Cortex. Current Biology, 23(15), 1427–1431. 10.1016/j.cub.2013.05.065

Amedi, A., Malach, R., & Pascual-Leone, A. (2005). Negative BOLD Differentiates Visual Imagery and Perception. Neuron, 48(5), 859–872. 10.1016/j.neuron.2005.10.032

Bae, G., & Luck, S. J. (2019). What happens to an individual visual working memory representation when it is interrupted? British Journal of Psychology, 110(2), 268–287. 10.1111/bjop.12339

Bainbridge, W. A., Pounder, Z., Eardley, A. F., & Baker, C. I. (2021). Quantifying aphantasia through drawing: Those without visual imagery show deficits in object but not spatial memory. Cortex, 135, 159–172. 10.1016/j.cortex.2020.11.014

Barbieri, R., Töpfer, F. M., Soch, J., Bogler, C., Sprekeler, H., & Haynes, J.-D. (2023). *Encoding of continuous perceptual choices in human early visual cortex* [Preprint]. bioRxiv. 10.1101/2023.02.10.527876

Bartolomeo, P. (2008). The neural correlates of visual mental imagery: An ongoing debate. Cortex, 44(2), 107–108. 10.1016/j.cortex.2006.07.001

Bartolomeo, P., Bachoud-Lévi, A.-C., De Gelder, B., Denes, G., Dalla Barba, G., Brugières, P., & Degos, J.-D. (1998). Multiple-domain dissociation between impaired visual perception and preserved mental imagery in a patient with bilateral extrastriate lesions. Neuropsychologia, 36(3), 239–249. 10.1016/S0028-3932(97)00103-6

Bartolomeo, P., Hajhajate, D., Liu, J., & Spagna, A. (2020). Assessing the causal role of early visual areas in visual mental imagery. Nature Reviews Neuroscience, 21, 2. 10.1038/s41583-020-0348-5

Bergmann, J., Genç, E., Kohler, A., Singer, W., & Pearson, J. (2016). Smaller Primary Visual Cortex Is Associated with Stronger, but Less Precise Mental Imagery. Cerebral Cortex, 26(9), 3838–3850. 10.1093/cercor/bhv186

Bettencourt, K. C., & Xu, Y. (2016). Decoding the content of visual short-term memory under distraction in occipital and parietal areas. Nature Neuroscience, 19(1), 150–157. 10.1038/nn.4174

Blajenkova, O., Kozhevnikov, M., & Motes, M. A. (2006). Object-spatial imagery: A new self-report imagery questionnaire. Applied Cognitive Psychology, 20(2), 239–263. 10.1002/acp.1182

Brainard, D. H. (1997). The Psychophysics Toolbox. Spatial Vision, 10(4), 433–436. 10.1163/156856897X00357

Brodersen, K. H., Ong, C. S., Stephan, K. E., & Buhmann, J. M. (2010). The Balanced Accuracy and Its Posterior Distribution. 2010 20th International Conference on Pattern Recognition, 3121–3124. 10.1109/ICPR.2010.764

Bullmore, E. T., Suckling, J., Overmeyer, S., Rabe-Hesketh, S., Taylor, E., & Brammer, M. J. (1999). Global, voxel, and cluster tests, by theory and permutation, for a difference between two groups of structural MR images of the brain. IEEE Transactions on Medical Imaging, 18(1), 32–42. 10.1109/42.750253

Chang, C.-C., & Lin, C.-J. (2011). LIBSVM: A library for support vector machines. ACM Transactions on Intelligent Systems and Technology, 2(3), 1–27. 10.1145/1961189.1961199

Chatterjee, A., & Southwood, M. H. (1995). Cortical blindness and visual imagery. Neurology, 45(12), 2189–2195. 10.1212/WNL.45.12.2189

Christophel, T. B., & Haynes, J.-D. (2014). Decoding complex flow-field patterns in visual working memory. NeuroImage, 91, 43–51. 10.1016/j.neuroimage.2014.01.025

Christophel, T. B., Hebart, M. N., & Haynes, J.-D. (2012). Decoding the Contents of Visual Short-Term Memory from Human Visual and Parietal Cortex. The Journal of Neuroscience, 32(38), 12983–12989. 10.1523/JNEUROSCI.0184-12.2012

Christophel, T. B., Klink, P. C., Spitzer, B., Roelfsema, P. R., & Haynes, J.-D. (2017). The Distributed Nature of Working Memory. Trends in Cognitive Sciences, 21(2), 111–124. 10.1016/j.tics.2016.12.007

Cichy, R. M., Heinzle, J., & Haynes, J.-D. (2012). Imagery and Perception Share Cortical Representations of Content and Location. Cerebral Cortex, 22(2), 372–380. 10.1093/cercor/bhr106

Cui, X., Jeter, C. B., Yang, D., Montague, P. R., & Eagleman, D. M. (2007). Vividness of mental imagery: Individual variability can be measured objectively. Vision Research, 47(4), 474–478. 10.1016/j.visres.2006.11.013

de Gelder, B., Tamietto, M., Pegna, A. J., & Van den Stock, J. (2015). Visual imagery influences brain responses to visual stimulation in bilateral cortical blindness. Cortex, 72, 15–26. 10.1016/j.cortex.2014.11.009

D’Esposito, M., & Postle, B. R. (2015). The Cognitive Neuroscience of Working Memory. Annual Review of Psychology, 66(1), 115–142. 10.1146/annurev-psych-010814-015031

Dijkstra, N., Bosch, S. E., & van Gerven, M. A. J. (2017). Vividness of Visual Imagery Depends on the Neural Overlap with Perception in Visual Areas. The Journal of Neuroscience, 37(5), 1367–1373. 10.1523/JNEUROSCI.3022-16.2016

Dijkstra, N., Bosch, S. E., & van Gerven, M. A. J. (2019). Shared Neural Mechanisms of Visual Perception and Imagery. Trends in Cognitive Sciences, 23(5), 423–434. 10.1016/j.tics.2019.02.004

Dijkstra, N., Mostert, P., Lange, F. P. de, Bosch, S., & van Gerven, M. A. (2018). Differential temporal dynamics during visual imagery and perception. ELife, 7, e33904. 10.7554/eLife.33904

Dijkstra, N., Zeidman, P., Ondobaka, S., van Gerven, M. A. J., & Friston, K. (2017). Distinct Top-down and Bottom-up Brain Connectivity During Visual Perception and Imagery. Scientific Reports, 7(1), 5677. 10.1038/s41598-017-05888-8

Drucker, H., Burges, C. J. C., Kaufman, L., Smola, A. J., & Vapnik, V. (1996). Support Vector Regression Machines. Advances in Neural Information Processing Systems, 9.

Ester, E. F., Anderson, D. E., Serences, J. T., & Awh, E. (2013). A Neural Measure of Precision in Visual Working Memory. Journal of Cognitive Neuroscience, 25(5), 754–761. 10.1162/jocn_a_00357

Ester, E. F., Serences, J. T., & Awh, E. (2009). Spatially Global Representations in Human Primary Visual Cortex during Working Memory Maintenance. Journal of Neuroscience, 29(48), 15258–15265. 10.1523/JNEUROSCI.4388-09.2009

Gazzaley, A., & Nobre, A. C. (2012). Top-down modulation: Bridging selective attention and working memory. Trends in Cognitive Sciences, 16(2), 129–135. 10.1016/j.tics.2011.11.014

Groppe, D. M., Urbach, T. P., & Kutas, M. (2011). Mass univariate analysis of event-related brain potentials/fields I: A critical tutorial review. Psychophysiology, 48(12), 1711–1725. 10.1111/j.1469-8986.2011.01273.x

Hallenbeck, G. E., Sprague, T. C., Rahmati, M., Sreenivasan, K. K., & Curtis, C. E. (2021). Working memory representations in visual cortex mediate distraction effects. Nature Communications, 12(1), 4714. 10.1038/s41467-021-24973-1

Harrison, S. A., & Tong, F. (2009). Decoding reveals the contents of visual working memory in early visual areas. Nature, 458(7238), 632–635. 10.1038/nature07832

Haynes, J.-D., & Rees, G. (2005). Predicting the orientation of invisible stimuli from activity in human primary visual cortex. Nature Neuroscience, 8(5), 686–691. 10.1038/nn1445

Hebart, M. N., Görgen, K., & Haynes, J.-D. (2015). The Decoding Toolbox (TDT): A versatile software package for multivariate analyses of functional imaging data. Frontiers in Neuroinformatics, 8, 88. 10.3389/fninf.2014.00088

Hubel, D. H., & Wiesel, T. N. (1968). Receptive fields and functional architecture of monkey striate cortex. The Journal of Physiology, 195(1), 215–243. 10.1113/jphysiol.1968.sp008455

Iamshchinina, P., Christophel, T. B., Gayet, S., & Rademaker, R. L. (2021). Essential considerations for exploring visual working memory storage in the human brain. Visual Cognition, 29(7), 425–436. 10.1080/13506285.2021.1915902

Jacobs, C., Schwarzkopf, D. S., & Silvanto, J. (2018). Visual working memory performance in aphantasia. Cortex, 105, 61–73. 10.1016/j.cortex.2017.10.014

Jeffreys, H. (1998). The theory of probability. OUP Oxford.

Kaas, A., Weigelt, S., Roebroeck, A., Kohler, A., & Muckli, L. (2010). Imagery of a moving object: The role of occipital cortex and human MT/V5+. NeuroImage, 49(1), 794–804. 10.1016/j.neuroimage.2009.07.055

Kamitani, Y., & Tong, F. (2005). Decoding the visual and subjective contents of the human brain. Nature Neuroscience, 8(5), 679–685. 10.1038/nn1444

Kay, L., Keogh, R., Andrillon, T., & Pearson, J. (2022). The pupillary light response as a physiological index of aphantasia, sensory and phenomenological imagery strength. ELife, 11, e72484. 10.7554/eLife.72484

Keogh, R., Bergmann, J., & Pearson, J. (2020). Cortical excitability controls the strength of mental imagery. ELife, 9, e50232. 10.7554/eLife.50232

Keogh, R., & Pearson, J. (2011). Mental Imagery and Visual Working Memory. PLoS ONE, 6(12), e29221. 10.1371/journal.pone.0029221

Keogh, R., & Pearson, J. (2014). The sensory strength of voluntary visual imagery predicts visual working memory capacity. Journal of Vision, 14(12), 7–7. 10.1167/14.12.7

Keogh, R., & Pearson, J. (2018). The blind mind: No sensory visual imagery in aphantasia. Cortex, 105, 53–60. 10.1016/j.cortex.2017.10.012

Keogh, R., Wicken, M., & Pearson, J. (2021). Visual working memory in aphantasia: Retained accuracy and capacity with a different strategy. Cortex, 143, 237–253. 10.1016/j.cortex.2021.07.012

Keysers, C., Gazzola, V., & Wagenmakers, E.-J. (2020). Using Bayes factor hypothesis testing in neuroscience to establish evidence of absence. Nature Neuroscience, 23(7), 788– 799. 10.1038/s41593-020-0660-4

Klein, I., Dubois, J., Mangin, J.-F., Kherif, F., Flandin, G., Poline, J.-B., Denis, M., Kosslyn, S. M., & Le Bihan, D. (2004). Retinotopic organization of visual mental images as revealed by functional magnetic resonance imaging. Cognitive Brain Research, 22(1), 26–31. 10.1016/j.cogbrainres.2004.07.006

Kleiner, M., Brainard, D., & Pelli, D. (2007). What’s new in Psychtoolbox-3? Perception, 36.

Kosslyn, S. M., Ganis, G., & Thompson, W. L. (2001). Neural foundations of imagery. Nature Reviews Neuroscience, 2(9), 635–642. 10.1038/35090055

Kosslyn, S. M., & Thompson, W. L. (2003). When is early visual cortex activated during visual mental imagery? Psychological Bulletin, 129(5), 723–746. 10.1037/0033-2909.129.5.723

Lamme, V. A. F., & Roelfsema, P. R. (2000). The distinct modes of vision offered by feedforward and recurrent processing. Trends in Neurosciences, 23(11), 571–579. 10.1016/S0166-2236(00)01657-X

Lee, S.-H., & Baker, C. I. (2016). Multi-Voxel Decoding and the Topography of Maintained Information During Visual Working Memory. Frontiers in Systems Neuroscience, 10. 10.3389/fnsys.2016.00002

Lee, S.-H., Kravitz, D. J., & Baker, C. I. (2012). Disentangling visual imagery and perception of real-world objects. NeuroImage, 59(4), 4064–4073. 10.1016/j.neuroimage.2011.10.055

Lee, S.-H., Kravitz, D. J., & Baker, C. I. (2013). Goal-dependent dissociation of visual and prefrontal cortices during working memory. Nature Neuroscience, 16(8), 997–999. 10.1038/nn.3452

Liu, T. (2016). Neural representation of object-specific attentional priority. NeuroImage, 129, 15–24. 10.1016/j.neuroimage.2016.01.034

Logie, R. H., Pernet, C. R., Buonocore, A., & Sala, S. D. (2011). Low and high imagers activate networks differentially in mental rotation. Neuropsychologia, 49(11), 3071–3077. 10.1016/j.neuropsychologia.2011.07.011

Love, J., Selker, R., Marsman, M., Jamil, T., Dropmann, D., Verhagen, J., Ly, A., Gronau, Q. F., Smíra, M., Epskamp, S., Matzke, D., Wild, A., Knight, P., Rouder, J. N., Morey, R. D., & Wagenmakers, E.-J. (2019). JASP: Graphical Statistical Software for Common Statistical Designs. Journal of Statistical Software, 88(2). 10.18637/jss.v088.i02

Maris, E., & Oostenveld, R. (2007). Nonparametric statistical testing of EEG- and MEG-data. Journal of Neuroscience Methods, 164(1), 177–190. 10.1016/j.jneumeth.2007.03.024

Marks, D. F. (1973). Visual Imagery Differences In The Recall Of Pictures. British Journal of Psychology, 64(1), 17–24. 10.1111/j.2044-8295.1973.tb01322.x

Mechelli, A. (2004). Where Bottom-up Meets Top-down: Neuronal Interactions during Perception and Imagery. Cerebral Cortex, 14(11), 1256–1265. 10.1093/cercor/bhh087

Merriam, E. P., Gardner, J. L., Movshon, J. A., & Heeger, D. J. (2013). Modulation of Visual Responses by Gaze Direction in Human Visual Cortex. Journal of Neuroscience, 33(24), 9879–9889. 10.1523/JNEUROSCI.0500-12.2013

Mongillo, G., Barak, O., & Tsodyks, M. (2008). Synaptic Theory of Working Memory. Science, 319(5869), 1543–1546. 10.1126/science.1150769

Morey, R. D., & Rouder, J. N. (2011). Bayes factor approaches for testing interval null hypotheses. Psychological Methods, 16(4), 406–419. 10.1037/a0024377

Moro, V., Berlucchi, G., Lerch, J., Tomaiuolo, F., & Aglioti, S. M. (2008). Selective deficit of mental visual imagery with intact primary visual cortex and visual perception. Cortex, 44(2), 109–118. 10.1016/j.cortex.2006.06.004

Naselaris, T., Olman, C. A., Stansbury, D. E., Ugurbil, K., & Gallant, J. L. (2015). A voxel-wise encoding model for early visual areas decodes mental images of remembered scenes. NeuroImage, 105, 215–228. 10.1016/j.neuroimage.2014.10.018

Oostenveld, R., Fries, P., Maris, E., & Schoffelen, J.-M. (2011). FieldTrip: Open Source Software for Advanced Analysis of MEG, EEG, and Invasive Electrophysiological Data. Computational Intelligence and Neuroscience, 2011, 1–9. 10.1155/2011/156869

Pearson, J. (2019). The human imagination: The cognitive neuroscience of visual mental imagery. Nature Reviews Neuroscience, 20(10), 624–634. 10.1038/s41583-019-0202-9

Pearson, J. (2020). Reply to: Assessing the causal role of early visual areas in visual mental imagery. Nature Reviews Neuroscience, 21, 2. 10.1038/s41583-020-0349-4

Pearson, J., Clifford, C. W. G., & Tong, F. (2008). The Functional Impact of Mental Imagery on Conscious Perception. Current Biology, 18(13), 982–986. 10.1016/j.cub.2008.05.048

Pearson, J., & Keogh, R. (2019). Redefining Visual Working Memory: A Cognitive-Strategy, Brain-Region Approach. Current Directions in Psychological Science, 28(3), 266–273. 10.1177/0963721419835210

Pearson, J., & Kosslyn, S. M. (2015). The heterogeneity of mental representation: Ending the imagery debate. Proceedings of the National Academy of Sciences, 112(33), 10089– 10092. 10.1073/pnas.1504933112

Pearson, J., Rademaker, R. L., & Tong, F. (2011). Evaluating the Mind’s Eye: The Metacognition of Visual Imagery. Psychological Science, 22(12), 1535–1542. 10.1177/0956797611417134

Pilly, P. K., & Seitz, A. R. (2009). What a difference a parameter makes: A psychophysical comparison of random dot motion algorithms. Vision Research, 49(13), 1599–1612. 10.1016/j.visres.2009.03.019

Purdon, P. L., & Weisskoff, R. M. (1998). Effect of temporal autocorrelation due to physiological noise and stimulus paradigm on voxel-level false-positive rates in fMRI. Human Brain Mapping, 6(4), 239–249. 10.1002/(SICI)1097-0193(1998)6:4<239::AID-HBM4>3.0.CO;2-4

Rademaker, R. L., & Pearson, J. (2012). Training Visual Imagery: Improvements of Metacognition, but not Imagery Strength. Frontiers in Psychology, 3. 10.3389/fpsyg.2012.00224

Ragni, F., Tucciarelli, R., Andersson, P., & Lingnau, A. (2020). Decoding stimulus identity in occipital, parietal and inferotemporal cortices during visual mental imagery. Cortex, 127, 371–387. 10.1016/j.cortex.2020.02.020

Reddy, L., Tsuchiya, N., & Serre, T. (2010). Reading the mind’s eye: Decoding category information during mental imagery. NeuroImage, 50(2), 818–825. 10.1016/j.neuroimage.2009.11.084

Rose, N. S., LaRocque, J. J., Riggall, A. C., Gosseries, O., Starrett, M. J., Meyering, E. E., & Postle, B. R. (2016). Reactivation of latent working memories with transcranial magnetic stimulation. Science, 354(6316), 1136–1139. 10.1126/science.aah7011

Senden, M., Emmerling, T. C., van Hoof, R., Frost, M. A., & Goebel, R. (2019). Reconstructing imagined letters from early visual cortex reveals tight topographic correspondence between visual mental imagery and perception. Brain Structure and Function, 224(3), 1167–1183. 10.1007/s00429-019-01828-6

Serences, J., Saproo, S., Scolari, M., Ho, T., & Muftuler, L. (2009). Estimating the influence of attention on population codes in human visual cortex using voxel-based tuning functions. NeuroImage, 44(1), 223–231. 10.1016/j.neuroimage.2008.07.043

Serences, J. T. (2016). Neural mechanisms of information storage in visual short-term memory. Vision Research, 128, 53–67. 10.1016/j.visres.2016.09.010

Serences, J. T., Ester, E. F., Vogel, E. K., & Awh, E. (2009). Stimulus-Specific Delay Activity in Human Primary Visual Cortex. Psychological Science, 20(2), 207–214. 10.1111/j.1467-9280.2009.02276.x

Spagna, A., Hajhajate, D., Liu, J., & Bartolomeo, P. (2021). Visual mental imagery engages the left fusiform gyrus, but not the early visual cortex: A meta-analysis of neuroimaging evidence. Neuroscience & Biobehavioral Reviews, 122, 201–217. 10.1016/j.neubiorev.2020.12.029

Sreenivasan, K. K., Curtis, C. E., & D’Esposito, M. (2014). Revisiting the role of persistent neural activity during working memory. Trends in Cognitive Sciences, 18(2), 82–89. 10.1016/j.tics.2013.12.001

Tanabe, J., Miller, D., Tregellas, J., Freedman, R., & Meyer, F. G. (2002). Comparison of Detrending Methods for Optimal fMRI Preprocessing. NeuroImage, 15(4), 902–907. 10.1006/nimg.2002.1053

Teng, C., & Kravitz, D. J. (2019). Visual working memory directly alters perception. Nature Human Behaviour, 3(8), 827–836. 10.1038/s41562-019-0640-4

Thorudottir, S., Sigurdardottir, H. M., Rice, G. E., Kerry, S. J., Robotham, R. J., Leff, A. P., & Starrfelt, R. (2020). The Architect Who Lost the Ability to Imagine: The Cerebral Basis of Visual Imagery. Brain Sciences, 10(2), 59. 10.3390/brainsci10020059

Tong, F. (2013). Imagery and visual working memory: One and the same? Trends in Cognitive Sciences, 17(10), 489–490. 10.1016/j.tics.2013.08.005

Töpfer, F. M., Barbieri, R., Sexton, C. M., Wang, X., Soch, J., Bogler, C., & Haynes, J.-D. (2022). Psychophysics and computational modeling of feature-continuous motion perception. Journal of Vision, 22(11), 16. 10.1167/jov.22.11.16

Ts’o, D. Y., Frostig, R. D., Lieke, E. E., & Grinvald, A. (1990). Functional Organization of Primate Visual Cortex Revealed by High Resolution Optical Imaging. Science, 249(4967), 417–420. 10.1126/science.2165630

Urai, A. E., Braun, A., & Donner, T. H. (2017). Pupil-linked arousal is driven by decision uncertainty and alters serial choice bias. Nature Communications, 8(1), 14637. 10.1038/ncomms14637

Wang, L., Mruczek, R. E. B., Arcaro, M. J., & Kastner, S. (2015). Probabilistic Maps of Visual Topography in Human Cortex. Cerebral Cortex, 25(10), 3911–3931. 10.1093/cercor/bhu277

Yun, K., Peng, Y., Samaras, D., Zelinsky, G. J., & Berg, T. L. (2013). Exploring the role of gaze behavior and object detection in scene understanding. Frontiers in Psychology, 4. 10.3389/fpsyg.2013.00917

Zeman, A., Dewar, M., & Della Sala, S. (2015). Lives without imagery – Congenital aphantasia. Cortex, 73, 378–380. 10.1016/j.cortex.2015.05.019

Zeman, A. Z. J., Della Sala, S., Torrens, L. A., Gountouna, V.-E., McGonigle, D. J., & Logie, R. H. (2010). Loss of imagery phenomenology with intact visuo-spatial task performance: A case of ‘blind imagination.’ Neuropsychologia, 48(1), 145–155. 10.1016/j.neuropsychologia.2009.08.024

Zhang, W., & Luck, S. J. (2008). Discrete fixed-resolution representations in visual working memory. Nature, 453(7192), 233–235. 10.1038/nature06860

